# Lapses in perceptual decisions reflect exploration

**DOI:** 10.1101/613828

**Authors:** Sashank Pisupati, Lital Chartarifsky-Lynn, Anup Khanal, Anne K. Churchland

## Abstract

Perceptual decision-makers often display a constant rate of errors independent of evidence strength. These “lapses” are treated as a nuisance arising from noise tangential to the decision, e.g. inattention or motor errors. Here, we use a multisensory decision task in rats to demonstrate that these explanations cannot account for lapses’ stimulus dependence. We propose a novel explanation: lapses reflect a strategic trade-off between exploiting known rewarding actions and exploring uncertain ones. We tested the model’s predictions by selectively manipulating one action’s reward magnitude or probability. As uniquely predicted by this model, changes were restricted to lapses associated with that action. Finally, we show that lapses are a powerful tool for assigning decision-related computations to neural structures based on disruption experiments (here, posterior striatum and secondary motor cortex). These results suggest that lapses reflect an integral component of decision-making and are informative about action values in normal and disrupted brain states.

## INTRODUCTION

Perceptual judgments are often modeled using noisy ideal observers (e.g., Signal detection theory, Green, Swets, et al., 1966; Bayesian decision theory, Dayan and Daw, 2008) that explain subjects’ errors as a consequence of noise in sensory evidence. This predicts an error rate that decreases with increasing sensory evidence, capturing the sigmoidal relationship often seen between evidence strength and subjects’ decision probabilities (i.e. the psychometric function; Fig. 1).

**Figure 1.**
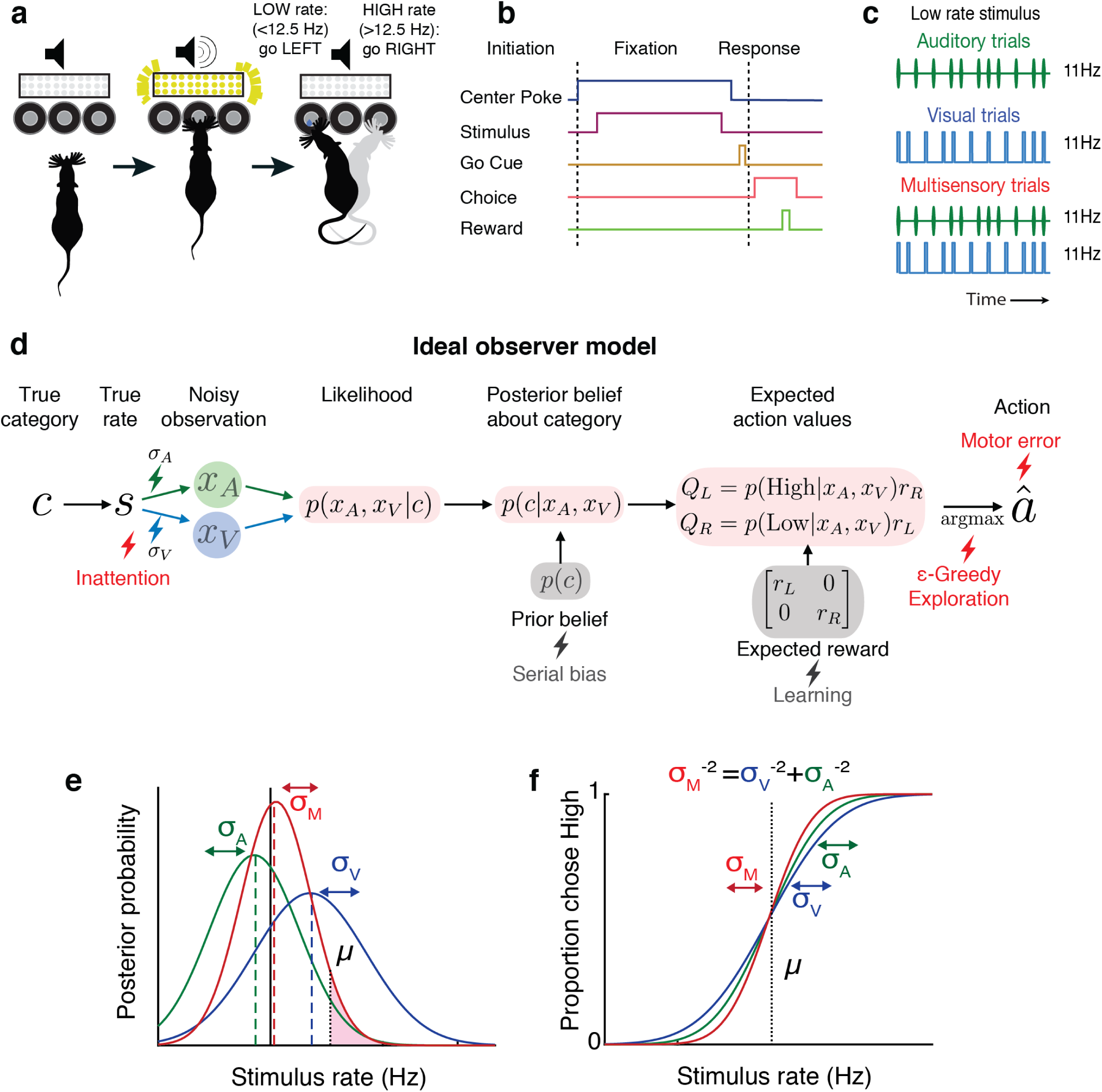
Testing ideal observer predictions in perceptual decision-making. **(a)** Schematic drawing of rate discrimination task. Rats initiate trials by poking into a center port. Trials consist of visual stimuli presented via a panel of diffused LEDs, auditory stimuli presented via a centrally positioned speaker or multisensory stimuli presented from both. Rats are rewarded with a 24*µ*l drop of water for reporting high rate stimuli (greater than 12.5 Hz) with rightward choices and low rate stimuli (lower than 12.5 Hz) with leftward choices. **(b)** Timeline of task events. **(c)** Example stimulus on auditory (top), visual (middle) and multisensory trials (bottom). Stimuli consist of a stream of events separated by long (100 ms) or short (50 ms) intervals. Multisensory stimuli consist of visual and auditory streams carrying the same underlying rate. Visual, auditory and multisensory trials were randomly interleaved (40% visual, 40% auditory, 20% multisensory). **(d)** Schematic outlining the computations of a Bayesian ideal observer. Stimulus belonging to a true category *c*, with a true underlying rate *s* gives rise to noisy observations *x*_*A*_ and *x*_*V*_, which are then integrated with each other and with prior beliefs to form a multisensory posterior belief about the category, and further combined with reward information to form expected action values *Q*_*L*_, *Q*_*R*_. The ideal observer selects the action *â* with maximum expected value. Lightning bolts denote proposed sources of noise that can give rise to (red) or exacerbate (grey) lapses, causing deviations from the ideal observer. **(e)** Posterior beliefs on an example trial assuming flat priors. Solid black line denotes true rate, blue and green dotted lines denote noisy visual and auditory observations, with corresponding unisensory posteriors shown in solid blue and green. Solid red denotes the multisensory posterior, centered around the M.A.P. rate estimate in dotted red. Shaded fraction denotes the probability of the correct choice being rightward, with *µ* denoting the category boundary. **(f)** Ideal observer predictions for the psychometric curve, i.e. proportion of high rate choices for each rate. Inverse slopes of the curves in each condition are reflective of the posterior widths on those conditions, assuming flat priors. The value on the abscissa corresponding to the curve’s midpoint indicates the subjective category boundary, assuming equal rewards and flat priors.

Human and non-human subjects often deviate from these predictions, displaying an additional constant rate of errors independent of the evidence strength known as “lapses”, leading to errors even on extreme stimulus levels (Wichmann and Hill, 2001; Busse et al., 2011; Gold and Ding, 2013; Carandini and Churchland, 2013). Despite the knowledge that ignoring or improperly fitting lapses can lead to serious mis-estimation of psychometric parameters (Wichmann and Hill, 2001; Prins and Kingdom, 2018), the cognitive mechanisms underlying lapses remain poorly understood. A number of possible sources of noise have been proposed to explain lapses, typically tangential to the decision-making process.

One class of explanations for lapses relies on pre-decision noise added due to fluctuating attention, which is often operationalized as a small fraction of trials on which the subject fails to attend to the stimulus (Wichmann and Hill, 2001). On these trials, it is assumed that the subject cannot specify the stimulus (i.e. sensory noise with infinite variance, Bays, Catalao, and Husain, 2009) and hence guesses randomly or in proportion to prior beliefs. This model can be thought of as a limiting case of the Variable Precision model, which assumes that fluctuating attention has a more graded effect of scaling the sensory noise variance (Garrido, Dolan, and Sahani, 2011), giving rise to heavy tailed estimate distributions, resembling lapses in the limit of high variability (Shen and Ma, 2019; Zhou et al., 2018). Temporal forms of inattention have also been proposed to give rise to lapses, where the animal ignores early or late parts of the evidence (impulsive or leaky integration, Erlich et al., 2015).

An alternative class of explanations for lapses relies on a fixed amount of noise added after a decision has been made, commonly referred to as “post-categorization” noise (Erlich et al., 2015) or decision noise (Law and Gold, 2009). Such noise could arise from errors in motor execution (e.g. finger errors, Wichmann and Hill, 2001), non-stationarities in the decision rule arising from computational imprecision (Findling et al., 2018), suboptimal weighting of choice or outcome history (Roy et al., 2018; Busse et al., 2011) or random variability added for the purpose of exploration (eg.“*ϵ*-greedy” decision rules).

A number of recent observations have cast doubt on fixed early- or late-stage noise as satisfactory explanations for lapses. For instance, many of these explanations predict that lapses should occur at a constant rate, while in reality, lapses are known to reduce in frequency with training in non-human primates (Law and Gold, 2009; Cloherty et al., 2019). Further, they can occur with different frequencies for different stimuli even within the same subject (in rodents, Nikbakht et al., 2018; and humans, Mihali et al., 2018; Bertolini et al., 2015; Flesch et al., 2018), suggesting that they may reflect task-specific, associative processes that can vary within a subject.

Lapse frequencies are even more variable across subjects and can depend on the subject’s age and state of brain function. For instance, lapses are significantly higher in children and patient populations than in healthy adult humans (Roach, Edwards, and Hogben, 2004; Witton, Talcott, and Henning, 2017; Manning et al., 2018). Moreover, a number of recent studies in rodents have found that perturbing neural activity in secondary motor cortex (Erlich et al., 2015) and striatum (Yartsev et al., 2018; Guo et al., 2018) has dramatic, asymmetric effects on lapses in auditory decision-making tasks. Because these perturbations were made in structures known to be involved in action selection, an intriguing possibility is that lapses reflect an integral part of the decision-making process, rather than a peripheral source of noise. However, because these studies only tested auditory stimuli, they did not afford the opportunity to distinguish sensory modality-specific deficits from general decision-related deficits. Taken together, these observations point to the need for a deeper understanding of lapses that accounts for effects of stimulus set, learning, age and neural perturbations.

Here, we leverage a multisensory decision-making task in rodents to reveal the inadequacy of traditional models in accounting for the variability of lapses across stimulus conditions of varying uncertainty. We re-examine a key assumption of perceptual decision-making theories, i.e. subject’s perfect knowledge of expected rewards (Dayan and Daw, 2008), to uncover a novel explanation for lapses: uncertainty-guided exploration, a well known strategy for balancing exploration and exploitation in value-based decisions. We confirm the predictions of the exploration model by manipulating the magnitude and probability of reward under conditions of varying uncertainty. Finally, we demonstrate that suppressing secondary motor cortex or posterior striatum unilaterally has an asymmetric effect on lapses that generalizes across sensory modalities, but only in uncertain conditions. This can be accounted for by an action value deficit contralateral to the inactivated side, reconciling the proposed perceptual and value-related roles of these areas and suggesting that lapses are informative about the subjective values of actions, reflecting a core component of decision-making.

## RESULTS

### Testing ideal observer predictions in perceptual decision-making

We leveraged an established decision-making task (Raposo, Sheppard, et al., 2012; Raposo, Kaufman, and Churchland, 2014; Sheppard, Raposo, and Churchland, 2013; Licata et al., 2017) in which freely moving rats judge whether the fluctuating rate of a 1000 ms series of auditory clicks and/or visual flashes (rate range: 9 - 16 Hz) is high or low compared with an abstract category boundary of 12.5Hz (Fig. 1a - c). Using Bayesian decision theory, we constructed an ideal observer for our task that selects choices that maximize expected reward (See Methods: Modelling). To test whether behavior matches ideal observer predictions, we presented multisensory trials with matched visual and auditory rates (i.e., both modalities carried the same number of events/sec; Fig. 1c, bottom) interleaved with visual-only or auditory-only trials. This allowed us to separately estimate the sensory noise in the animals’ visual and auditory system, and compare the measured performance on multisensory trials to the predictions of the ideal observer.

Performance was assessed using a psychometric curve, i.e. the probability of high-rate decisions as a function of stimulus rate (Fig. 1f). The ideal observer model predicts a relationship between the slope of the psychometric curve and noise in the animal’s estimate: the higher the standard deviation (*σ*) of sensory noise, the more uncertain the animals estimate of the rate and the shallower the psychometric curve. On multisensory trials, the ideal observer should have a more certain estimate of the rate (Fig. 1e, visual [blue] and auditory [green] *σ* values are larger than multisensory *σ* [red]), driving a steeper psychometric curve (Fig. 1f, red curve is steeper than green and blue curves). Since this model does not take lapses into account, it would predict perfect performance on the easiest stimuli regardless of uncertainty, and thus all curves should asymptote at 0 and 1 (Fig 1f).

### Lapses cause deviations from ideal observer, and are reduced on multisensory trials

In practice, the shapes of empirically obtained psychometric curves do not perfectly match the ideal observer (Fig. 2) since they asymptote at values that are less than 1 or greater than 0. This is a well known phenomenon in psychophysics (Wichmann and Hill, 2001), requiring two additional lapse parameters to precisely capture the asymptotes. To account for lapses, we fit a four-parameter psychometric function to the subjects’ choice data (Fig. 2a - red, Equation 1 in Methods) with the Palamedes toolbox (Prins and Kingdom, 2018). *γ* and *λ* are the lower and upper asymptote of the psychometric function, which parameterize lapses on low and high rates respectively; *ϕ* is a sigmoidal function, in our case the cumulative normal distribution; *x* is the event rate, i.e. the average number of flashes or beeps presented during the one second stimulus period; *µ* parameterizes the midpoint of the psychometric function and *σ* describes the inverse slope after correcting for lapses.

**Figure 2.**
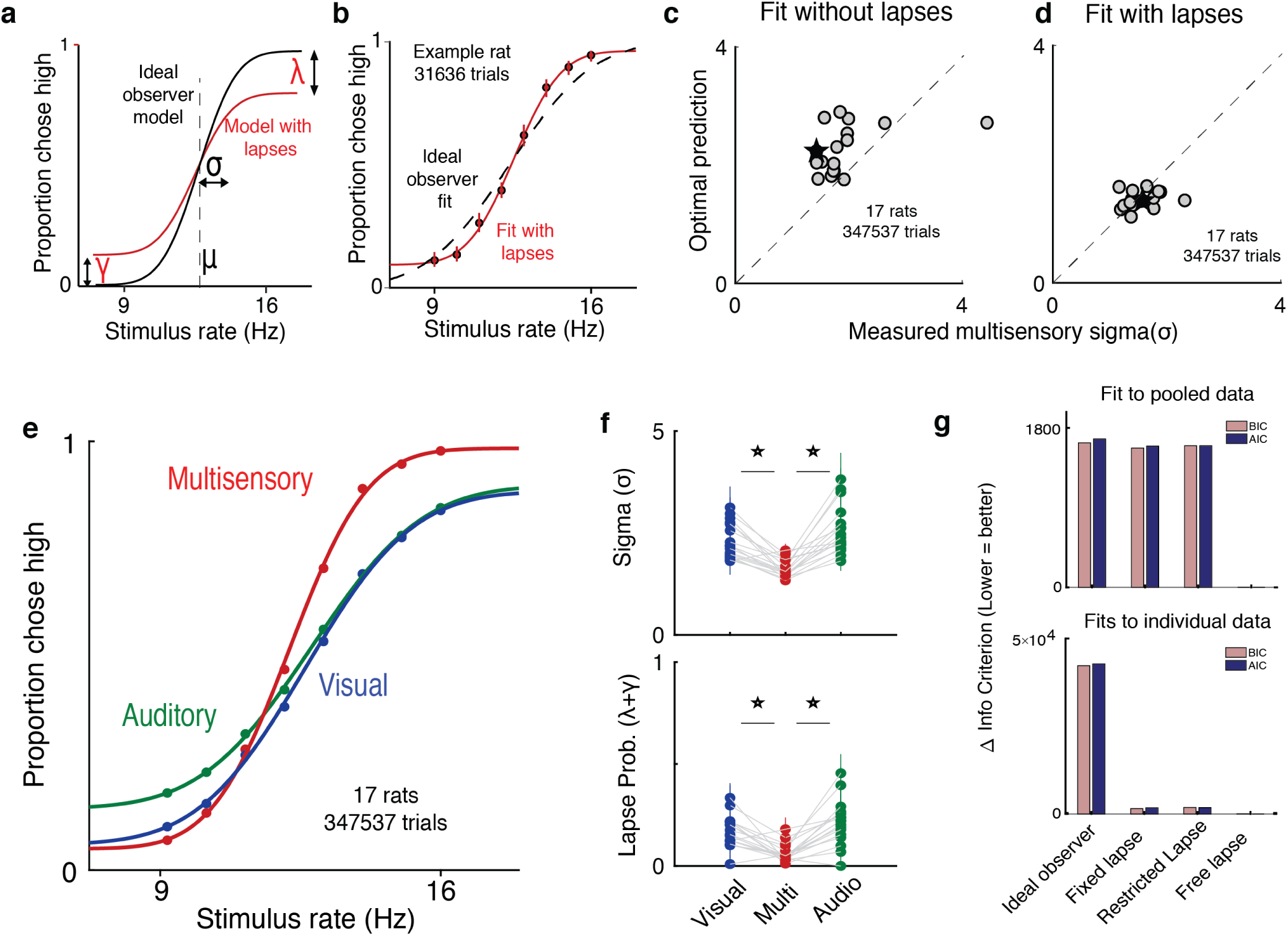
Deviations from ideal observer reflect lapses in judgment. **(a)** Schematic psychometric performance of an ideal observer (black) vs. a model that includes lapses (red). The ideal observer model includes two parameters: midpoint (*µ*) and inverse slope (*σ*). The four-parameter model includes *µ, σ*, and lapse probabilities for low rate (*γ*) and high rate choices (*λ*). Dotted line shows the true category boundary (12.5 Hz). **(b)** Subject data was fit with an ideal observer model (black) and a four-parameter model (red). **(c**,**d)** Ideal observer predictions vs. measured multisensory sigma for fits with and without lapses. **(c)** Multisensory integration seems supra-optimal when not accounting for lapses. **(d)** Optimal multisensory integration is restored when accounting for lapses. (n = 17 rats. Points represent individual rats; star represents pooled data across all rats. Data points that lie on the unity line represent cases in which the measured sigma was equal to the optimal prediction). **(e)** Rats’ psychometric curves on auditory (green), visual (blue) and multisensory (red) trials. Points represent data pooled across 17 rats, lines represent separate four-parameter fits to each condition. **(f)** Fit values of sigma (top) and lapse parameters (bottom) on unisensory and multisensory conditions. Both parameters showed significant reduction on the multisensory conditions (paired t-test, p<0.05); n=17 rats (347537 trials). **(g)** Model comparison using BIC (pink) and AIC (blue) for fits to pooled data across subjects (top) and to individual subject data (bottom). Lower scores indicate better fits. Both metrics favor a model where lapses are allowed to vary freely across conditions (“Free lapse”) over one without lapses (“Ideal observer”), one with a fixed probability of lapses (“Fixed lapse”) or where the lapses are restricted to being less than 0.1 (“Restricted lapse”).

How can we be sure that the asymptotes seen in the data truly reflect non-zero asymptotes rather than fitting artifacts or insufficient data at the asymptotes? To test whether lapses were truly necessary to explain the behavior, we fit the curves with and without lapses (Fig. 2b) and tested whether the lapse parameters were warranted. The fit without lapses was rejected in 15/17 rats by the Bayes Information Criterion (BIC), and in all rats by the Akaike Information Criterion (AIC). Fitting a fixed lapse rate across conditions was not sufficient to capture the data, nor was fitting a lapse rate that was constrained to be less than 0.1 (Wichmann and Hill, 2001). Both data pooled across subjects and individual subject data warranted fitting separate lapse rates to each condition (“free lapses” model outperforms “fixed lapses”, “restricted lapses” or “no lapses/ideal observer” in Fig. 2g).

Multisensory trials offer an additional, strong test of ideal observer predictions. In addition to perfect performance on the easiest stimuli, the ideal observer model predicts the minimum possible uncertainty achievable on multisensory trials through optimal integration (Ernst and Bulthoff, 2004; Equation 9 in Methods). By definition, better-than-optimal performance is impossible. However, studies in humans, rodents and non-human primates performing multisensory decision-making tasks suggest that in practice, performance occasionally exceeds optimal predictions (Raposo, Sheppard, et al., 2012; Nikbakht et al., 2018; Hou et al., 2018), seeming, at first, to violate the ideal observer model. Moreover, in these datasets, performance on the easiest stimuli was not perfect and asymptotes deviated from 0 and 1. As in these previous studies, when we fit performance without lapses, multisensory performance was significantly supra-optimal (p=0.0012, paired t-test), i.e. better than the ideal observer prediction (Fig. 2c, data points are above the unity line). This was also true when lapse probabilities were assumed to be fixed across conditions (p =0.0018) or when they were assumed to be less than 0.1 (p=0.0003). However, when we allowed lapses to vary freely across conditions, performance was indistinguishable from optimal (Fig. 2d, data points are on the unity line). This reaffirms that proper treatment of lapses is crucial for accurate estimation of perceptual parameters and offers a potential explanation for previous reports of supra-optimality.

Using this improved fitting method, we replicated previous observations (Raposo, Sheppard, et al., 2012; Raposo, Kaufman, and Churchland, 2014) showing that animals have improved sensitivity (lower *σ*) on multisensory vs. unisensory trials (Fig. 2e, red curve is steeper than green/blue curves; Fig. 2f, top). Interestingly, we observed that animals also had a lower lapse probability (*λ* + *γ*) on multisensory trials (Fig. 2e, asymptotes for red curve are closer to 0 and 1; n=17 rats, 347537 trials). This was consistently observed across animals (Fig. 2f bottom, the probability of lapses on multisensory trials was 0.06 on average, compared to 0.17 on visual, p=1.4e-4 and 0.21 on auditory, p=1.5e-5).

### A novel model, uncertainty-guided exploration, explains lapses better than traditional models of inattention or motor-error

What could account for the reduction in lapse probability on multisensory trials? While adding extra parameters to the ideal observer model fit the behavioral data well and accounted for the reduction in inverse-slope on multisensory trials, this success doesn’t provide an explanation for why lapses are present in the first place, nor why they differ between stimulus conditions.

To investigate this, we considered possible sources of noise that have traditionally been invoked to explain lapses (Fig. 1d). We first hypothesized that lapses might be due to a fixed amount of noise added once the decision has been made. These sources of noise could include decision noise due to imprecision (Findling et al., 2018) or motor errors (Wichmann and Hill, 2001). However, these sources should hinder decisions equally across stimulus conditions (Supplementary Fig. 1b), which cannot explain our observation of condition-dependent lapse rates (Fig. 2f).

A second explanation is that lapses arise due to inattention on a small fraction of trials. Inattention would drive the animal to guess randomly, producing lapse rates whose sum should reflect the probability of not attending (Fig. 3a, Methods). According to this explanation, the lower lapse rate on multisensory trials reflects increased attention on those trials, perhaps due to their increased bottom-up salience (i.e. two streams of stimuli instead of one). To test this, we leveraged a multisensory condition that has been used to manipulate uncertainty without changing salience in rats and humans (Raposo, Sheppard, et al., 2012). Specifically, we interleaved standard matched-rate multisensory trials with “neutral” multisensory trials for which the rate of the auditory stimuli ranged between 9-16 Hz, while the visual stimuli was always 12 Hz. This rate was so close to the category boundary (12.5 Hz) that it did not provide compelling evidence for one choice or the other (Fig. 3d, left), thus reducing the information in the multisensory stimulus and increasing uncertainty. However, since both “neutral” and “matched” conditions are multisensory, they should be equally salient, and since they are interleaved, the animal would be unable to identify the condition without actually attending to the stimulus. According to the inattention model, matched and neutral trials should have the same rate of lapses, only differing in their *σ* (Supplementary Fig 1c).

**Figure 3.**
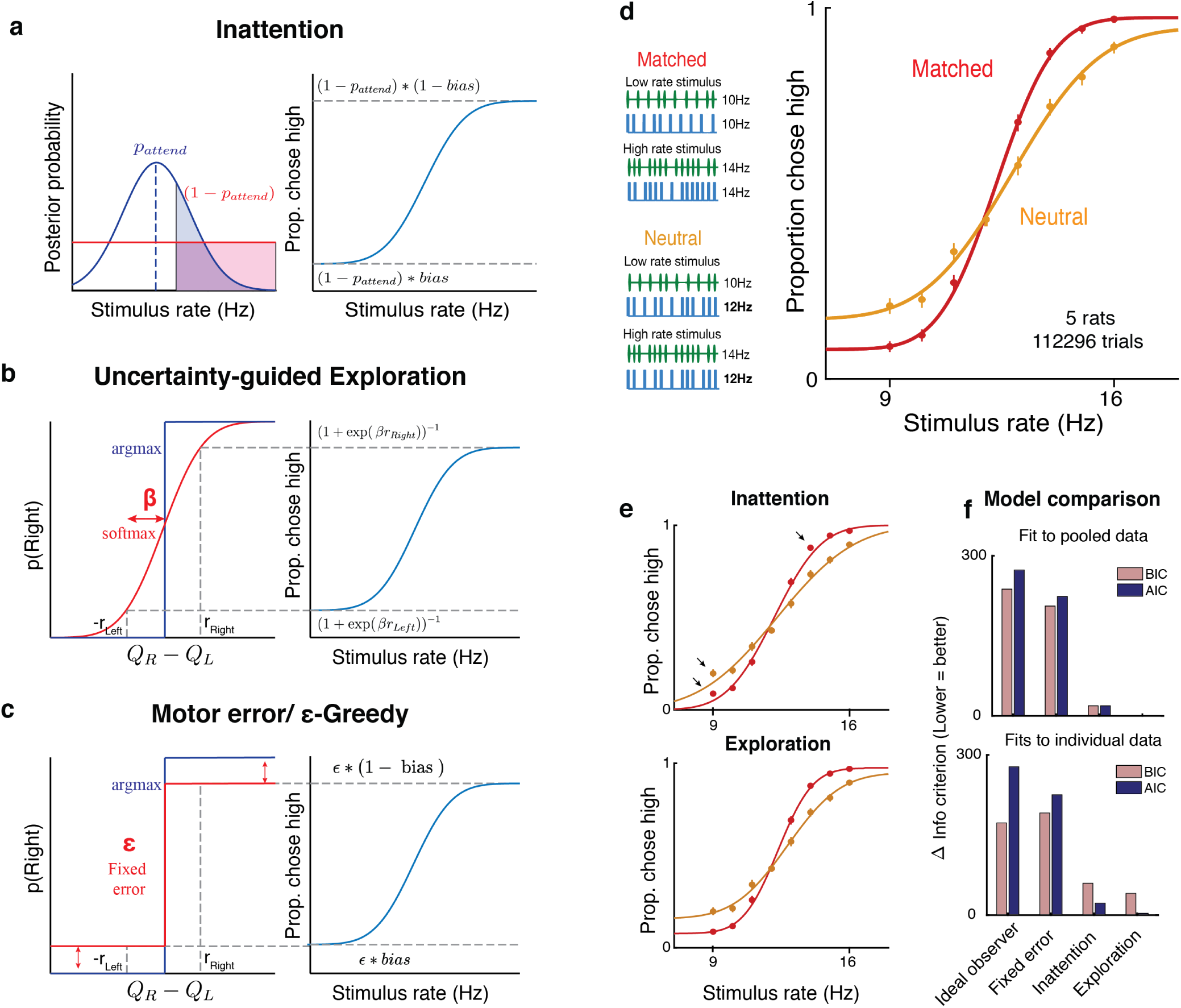
Uncertainty-guided exploration offers a novel explanation for lapses. Models of lapses in decision-making: **(a)** Inattention model of lapses. Left panel: on a small fraction of trials given by 1 −*p*_*attend*_, the observer does not attend to the stimulus (red curve), leading to equal posterior probabilities of high and low rates (Shaded, clear regions of curve respectively) and guesses according to the probability *bias*, giving rise to lapses (right panel). The sum of lapse rates then reflects 1 − *p*_*attend*_, while their ratio reflects the *bias*. **(b)** Uncertainty-guided exploration model. Lapses can arise from exploratory decision rules such as the “softmax” (red) rather than reward-maximization (blue). Since the difference in expected value from right and left actions (*Q*_*R*_ −*Q*_*L*_) is bounded by the maximum reward magnitudes *r*_*Right*_ and *r*_*Left*_, even when the stimulus is very easy, the maximum probability of choosing the higher value option is not 1, giving rise to lapses. Lapse rates on either side are then proportional to the reward magnitude on that side, and to a “temperature” parameter *β* that depends on the uncertainty in expected reward. **(c)** Motor error, or *ϵ*-greedy model. Lapses can also arise from decision rules with a fixed proportion *ϵ* of random exploratory choices, or due to motor errors ocurring on *ϵ* fraction of trials. The sum of lapses reflects *ϵ* while their ratio reflects any *bias* in exploration or motor errors. **(d)** Left: multisensory stimuli designed to distinguish between attentional and non-attentional sources of lapses. Standard multisensory stimuli with matched visual and auditory rates (top) and “neutral” stimuli where one modality has a rate very close to the category boundary and is uninformative (bottom). Both stimuli are multisensory and designed to have equal bottom-up salience, and can only be distinguished by attending to them and accumulating evidence. Right: rat performance on interleaved matched (red) and neutral (orange) trials. **(e)** Since the matched and neutral conditions are equally salient, they are expected to have equal probabilities of attending, predicting similar total lapse rates in the inattention model (top, solid lines are model fits). Deviations from model fits are denoted with arrows. The exploration model (bottom) provides a better fit, by allowing for different levels of exploration in the two conditions. **(f)** Model comparison using BIC (pink) and AIC (blue) both favor the uncertainty-guided exploration model for pooled data (top) as well as individual subject data (bottom).

Contrary to this prediction, we observed higher lapse rates on “neutral” trials, where the uncertainty was high, than on “matched” trials, where the uncertainty was low (Fig. 3d). The dependence of lapses on uncertainty is reminiscent of the dependence of lapses on uncertainty observed when comparing unisensory vs. multisensory trials (Fig. 2e,f; Supplementary Fig. 1e).

Having observed that traditional explanations of lapses fail to account for the behavioral observations, we re-examined a key assumption of ideal observer models used in perceptual decision-making - that subjects have complete knowledge about the rules and rewards (Dayan and Daw, 2008). In general, this assumption may not hold true for a number of reasons - even when the stimulus category is known, subjects might have uncertainty about the values of different actions because they are still in the process of learning (Law and Gold, 2009), because they incorrectly assume that their environment is non-stationary (Yu and Cohen, 2009), or because they forget over time (Gershman, 2015; Drugowitsch and Pouget, 2018). In such situations, rather than always “exploiting” (picking the action currently assumed to have the highest value), it is advantageous to “explore” (occasionally picking actions whose value the subject is uncertain about), in order to gather more information and maximize reward in the long term (Dayan and Daw, 2008). Exploratory choices of the lower value action for the easiest stimuli would resemble lapses, and the sum of lapses would reflect the overall degree of exploration.

Choosing how often to explore is challenging, and requires trading off immediate rewards for potential gains in information - random exploration would reward subjects at chance, but would reduce uncertainty uniformly about the value of all possible stimulus-action pairs, while a greedy policy (i.e. always exploiting) would yield many immediate rewards while leaving lower value stimulus-action pairs highly uncertain (Supplementary Fig. 2a,b). Policies that explore randomly on a small fraction of trials (e.g. “*ϵ*-Greedy” policies) do not make prescriptions about how often the subject should explore, and are behaviorally indistinguishable from motor errors when the fraction is fixed (Fig. 3c). One elegant way to automatically balance exploration and exploitation is to explore more often when one is more uncertain about action values. In particular, a form of uncertainty-guided exploration called Thompson sampling is known to be asymptotically optimal in many general environments (Leike et al., 2016), and reduces to a dynamic “softmax” decision rule (Fig. 3b), whose “inverse temperature” parameter (*β*) scales with uncertainty (Gershman, 2018). This predicts a lower *β* when values are more uncertain, encouraging more exploration and more frequent lapses, and a higher *β* when values are more certain, encouraging exploitation. The limiting case of perfect knowledge (*β* → *∞*) reduces to the reward-maximizing ideal observer.

The subjects’ task is further complicated by perceptual uncertainty - on trials where the stimulus category is not fully known, credit cannot be unambiguously assigned to one stimulus-action pair when rewards are obtained (Lak et al., 2018). This predicts that conditions with higher perceptual uncertainty (e.g. unisensory or neutral trials) should have more overlapping value beliefs (Supplementary Fig. 2c), encouraging more exploration and giving rise to more frequent lapses (Supplementary Fig. 2d).

As a result, on multisensory neutral trials, the uncertainty-guided exploration model predicts an increase not only in the *σ* parameter, but also in lapses, just as we observed (Fig. 3d). In fact, this model predicts that both these parameters should match those on auditory trials, since these conditions have comparable levels of perceptual uncertainty. This model fit the data well (Fig. 3e, bottom). By contrast, the inattention model predicts that both conditions would have the same lapse rates, with the neutral condition simply having a larger *σ*. This model provided a worse fit to the data, particularly missing the data at extreme stimulus values where lapses are most clearly apparent (Fig. 3e, top). Model comparison using BIC and AIC favored the exploration model over the inattention model, both for fits to pooled data across subjects (Fig. 3f top) and fits to individual subject data (Fig. 3f bottom, Supplementary Fig. 3)

### Reward manipulations confirm predictions of exploration model

One of the key features of the uncertainty-guided exploration model is that lapses are exploratory choices made with full knowledge of the stimulus, and should therefore depend only on the expected rewards associated with that stimulus category (Supplementary Fig. 2). This is in stark contrast to the inattention model and many other kinds of disengagement (Supplementary Fig. 4). According to these models, lapses are caused by the observer disregarding the stimulus, and hence lapses at the two extreme stimulus levels are influenced by a common underlying guessing process that depends on expected rewards from both stimulus categories. This is also in contrast to fixed motor error or *ϵ*-greedy models in which lapses are independent of expected reward (Fig. 3c).

Therefore, a unique prediction of the exploration model is that selectively manipulating expected rewards associated with one of the stimulus categories should only affect lapses at one extreme of the psychometric function. Conversely, inattention and other kinds of disengagement predict that both lapses should be affected, while fixed error models predict that neither should be affected (Fig. 4a, Supplementary Fig. 1,4).

**Figure 4.**
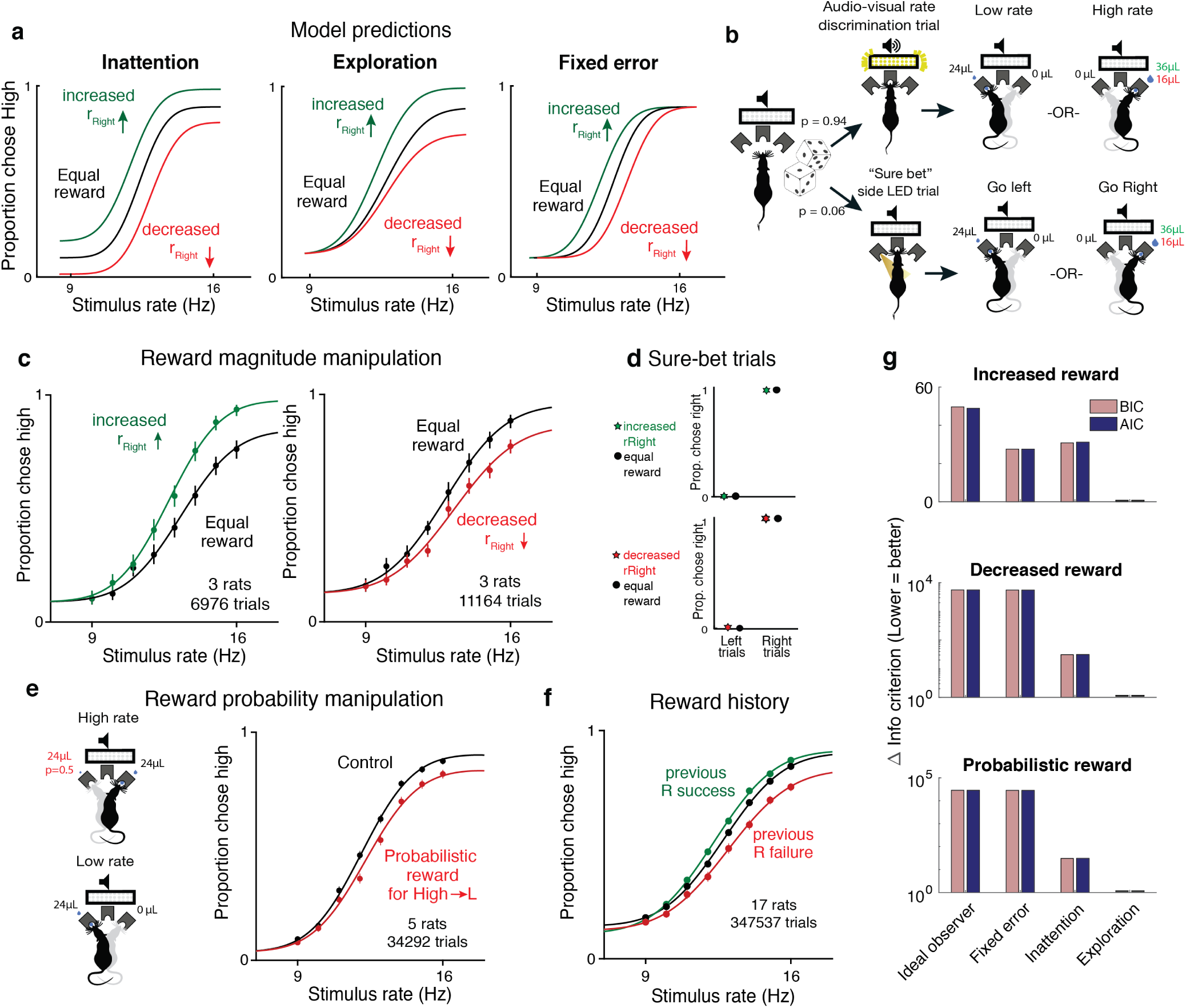
Reward manipulations match predictions of the exploration model. **(a)** The inattention, exploration and fixed error models make different predictions for increases and decreases in the reward magnitude for rightward (high-rate) actions. The inattention model (left panel) predicts changes in lapses for both high and low rate choices, while the exploration model (center panel) predicts changes in lapses only for high rate choices, and fixed motor error or *ϵ*-greedy models (right) predict changes in neither lapse. Black line, equal rewards on both sides; green, increased rightward reward; red, decreased rightward reward. **(b)** Schematic of rate discrimination trials and interleaved “sure bet” trials. The majority of the trials (94%) were rate discrimination trials as described in Figure 1. On sure-bet trials, a pure tone was played during a 0.2 second fixation period and one of the side ports was illuminated once the tone ended to indicate that reward was available there. Rate discrimination and sure-bet trials were randomly interleaved, as were left and right trials, and the rightward reward magnitude was either increased (36*µ*l) or decreased (16*µ*l) while maintaining the leftward reward at 24*µ*l **(c)** Rats’ behavior on rate discrimination trials following reward magnitude manipulations. High rate lapses decrease when water reward for high-rate choices is increased (left panel; n=3 rats, 6976 trials), while high-rate lapses increase when reward on that side is decreased (right panel; n=3 rats, 11164 trials). Solid curves are exploration model fits with a single parameter change accounting for the manipulation. **(d)** Rats show nearly perfect performance on sure-bet trials, and are unaffected by reward manipulations on these trials. **(e)** Reward probability manipulation. (Left) Schematic of probabilistic reward trials, incorrect (leftward) choices on high rates were rewarded with a probability of 0.5, and all other rewards were left unchanged. (Right) Rats’ behavior and exploration model fits showing a selective increase in high-rate lapses (n=5 rats, 34292 trials). **(f)** Rats’ behavior on equal reward trials conditioned on successes (green) or failures (red) on the right on the previous trials resembles effects of reward size manipulations. **(g)** Model comparison showing that AIC and BIC both favor the exploration model on data from all 3 manipulations.

To experimentally test these predictions, we tested rats on the rate discrimination task with asymmetric rewards (Fig. 4b, top). Instead of rewarding high and low rate choices equally, we increased the water amount on the reward port associated with high-rates (rightward choices) so it was 1.5 times larger than before, without changing the reward on the the low-rate side (leftward choices). In a second rat cohort we did the opposite: we devalued the choices associated with high-rate trials by decreasing the water amount on that side port so it was 1.5 times smaller than before, without changing the reward on the low-rate side.

The animals’ behavior on the asymmetric-reward task matched the predictions of the exploration model. Increasing the reward size on choices associated with high-rates led to a decrease in lapses for the highest rates and no changes in lapses for the lower rates (Fig. 4c, left; n=3 rats, 6976 trials). Decreasing the reward of choices associated with high-rates led to an increase in lapses for the highest rates and no changes in lapses for the lower rates (Fig. 4c, right; n=3 rats, 11164 trials). This shows that both increasing and decreasing the value of one of the actions has an asymmetric effect on lapse probabilities that does not match the inattention model.

To confirm that the asymmetric changes in lapse rate that we observed were truly driven by uncertainty, we examined performance on randomly interleaved “sure bet” trials on which the uncertainty was very low (Fig. 4b, bottom). On these trials, a pure tone was played during the fixation period, after which an LED at one of the side ports was clearly illuminated, indicating a reward. Sure-bet trials comprised 6% of the total trials, and as with the rate discrimination trials, left and right trials were interleaved. Owing to the low uncertainty, the model predicts that very little exploration would be required in this condition, and that animals would very quickly reach perfect performance on these trials. Importantly, our model predicts that performance on “sure-bet” trials would be unaffected by imbalances in reward magnitude.

In keeping with this prediction, performance on sure-bet trials was near perfect (rightward probabilities of 0.003 [0.001,0.01] and 0.989 [0.978,0.995] on go-left and go-right trials respectively), and unaffected following reward manipulations (Fig. 4d: Rightward probabilities of 0.004 [0.001, 0.014] and 0.996 [0.986,0.999] on increased reward, 0.006 [0.003,0.012] and 0.99 [0.983,0.994] on decreased reward). This suggests that the effect of action value on lapses is restricted to uncertain situations that encourage subjects to explore, rather than exploit.

As an additional test of the model, we manipulated expected rewards by probabilistically rewarding incorrect choices. Here, leftward choices on high rate (“go right”) trials were rewarded with a probability of 0.5, while leaving all other rewards unchanged (Fig. 4e left). The exploration model predicts that this should selectively increase the value of leftward actions on high rate trials, increasing lapses on high rates. Indeed, this is what we observed (Fig. 4e right, n=5 animals, 347537 trials), and the effect was strikingly similar to the decreased reward experiment, even though the two manipulations affect high rate action values through changes on opposite reward ports. Moreover, this suggests that lapses reflect changes in action value caused by changing either reward magnitudes or reward probabilities, as one would expect from the exploration model.

An added consequence of uncertainty in action values is that it should encourage continued learning even in the absence of explicit reward manipulations. This means that animals should continue to use the outcomes of previous trials to update the values of different actions as long as this uncertainty persists. Such persistent learning has been observed in a number of studies (Busse et al., 2011; Lak et al., 2018; Mendonca et al., 2018; Odoemene et al., 2018; Pinto et al., 2018; Scott et al., 2015). The uncertainty-dependent exploration model predicts that changes in action values should manifest as changes in lapse rates. For example, action value of rightward choices should increase following a rightward success and decrease following a rightward failure. This predicts similar changes in lapses as reward magnitude manipulations. As predicted, trials following rewarded and unrewarded rightward choices showed decreased and increased lapses, respectively (Fig. 4f; same rats and trials as in Fig. 2e). Taken together, manipulations of value confirm the predictions of the uncertainty-dependent exploration model (Fig. 4g).

### Lapses are a powerful tool for assigning decision-related computations to neural structures based on disruption experiments

The results of the behavioral manipulations (above) predict that unilateral disruption of areas that compute action values should affect lapse rates asymmetrically. In contrast, disruptions to areas that process sensory evidence would lead to horizontal biases without affecting lapses, and disruptions to motor areas that make one of the actions harder to perform would affect both lapses (Supplementary Fig. 5a). Crucially, in the absence of lapses, all three of these disruptions would drive an identical behavioral effect, a horizontal shift of the psychometric function (Supplementary Fig. 5b). This suggests that lapses can be used as a tool to determine which computations are affected by disruptions of a candidate brain region. To demonstrate this, we identified two candidate areas, secondary motor cortex (M2) and posterior striatum (pStr), that receive convergent input from primary visual and auditory cortices (Supplementary Fig. 6, results of simultaneous anterograde tracing from V1 and A1; also see Jiang and Kim, 2018; Barthas and Kwan, 2017). In previous work, disruptions of these areas had effects on auditory decisions, including changes in lapses (Erlich et al., 2015; Guo et al., 2018). However, considerable controversy remains as to which computations were affected by those disruptions. The effects were largely interpreted in terms of traditional ideal observer models, and thus attributed to perceptual biases (Guo et al., 2018), leaky accumulation (Erlich et al., 2015) or post categorization biases (Piet et al., 2017; Erlich et al., 2015). However, the asymmetric effects on lapses seen in these studies resembled the effects of the reward manipulations in our task, hinting that they may actually arise from action value changes. Importantly, these existing studies used only auditory stimuli, so were limited in their ability to distinguish sensory-specific deficits from action value deficits.

Here, we used analyses of lapses to determine the decision-related computations altered by unilateral disruption of M2 and pStr. If these disruptions affected action values, the exploration model makes three strong predictions. First, because action values are computed late in the decision-process, the model predicts that the effects should not depend on the modality of the stimulus. We therefore performed disruptions in animals doing interleaved auditory, visual and multisensory trials. If pStr and M2 indeed compute action value, then following unilateral disruption of these areas, our model should capture changes to all three modalities by a single parameter change to the contralateral action value. Second, these disruptions should selectively affect lapses on stimuli associated with contralateral actions, irrespective of the stimulus-response contingency. To test this, we performed disruptions on animals trained on standard and reversed contingencies. Finally, because altered action values should have no effect when there is no uncertainty and consequently no exploration, disruption to pStr and M2 should spare performance on sure-bet trials (Fig. 4b, bottom).

We suppressed activity of neurons in each of these areas using muscimol, a *GABA*_*A*_ agonist, during our multisensory rate discrimination task. We implanted bilateral cannulae in M2 (Fig. 5a, Supplementary Fig. 7b; n = 5 rats; +2 mm AP 1.3 mm ML, 0.3 mm DV) and pStr (Fig. 5a, Supplementary Fig. 7a; n = 6 rats; −3.2 mm AP, 5.4 mm ML, 4.1 mm DV). On control days, rats were infused unilaterally with saline, followed by unilateral muscimol infusion the next day (M2: 0.1-0.5 *µ*g, pStr 0.075-0.125 *µ*g). We compared performance on the multisensory rate discrimination task for muscimol days with preceding saline days. Inactivation of the side associated with low-rate choices biased the animals to make more low-rate choices (Fig. 5b; left 6 panels: empty circles, inactivation sessions; full circles, control sessions), while inactivation of the side associated with high-rates biased them to make more high-rate choices (Fig. 5b, right 6 panels). The inactivations largely affected lapses on the stimulus rates associated with contralateral actions, while sparing those associated with ipsilateral actions (Fig. 5c). These results recapitulated previous findings, and were strikingly similar to the effects we observed following reward manipulations (as seen in Fig. 4c, right panel). These effects were seen across areas (Fig. 5b, top, M2; bottom, pStr) and modalities (Fig. 5b; green, auditory; blue, visual and red, multisensory).

**Figure 5.**
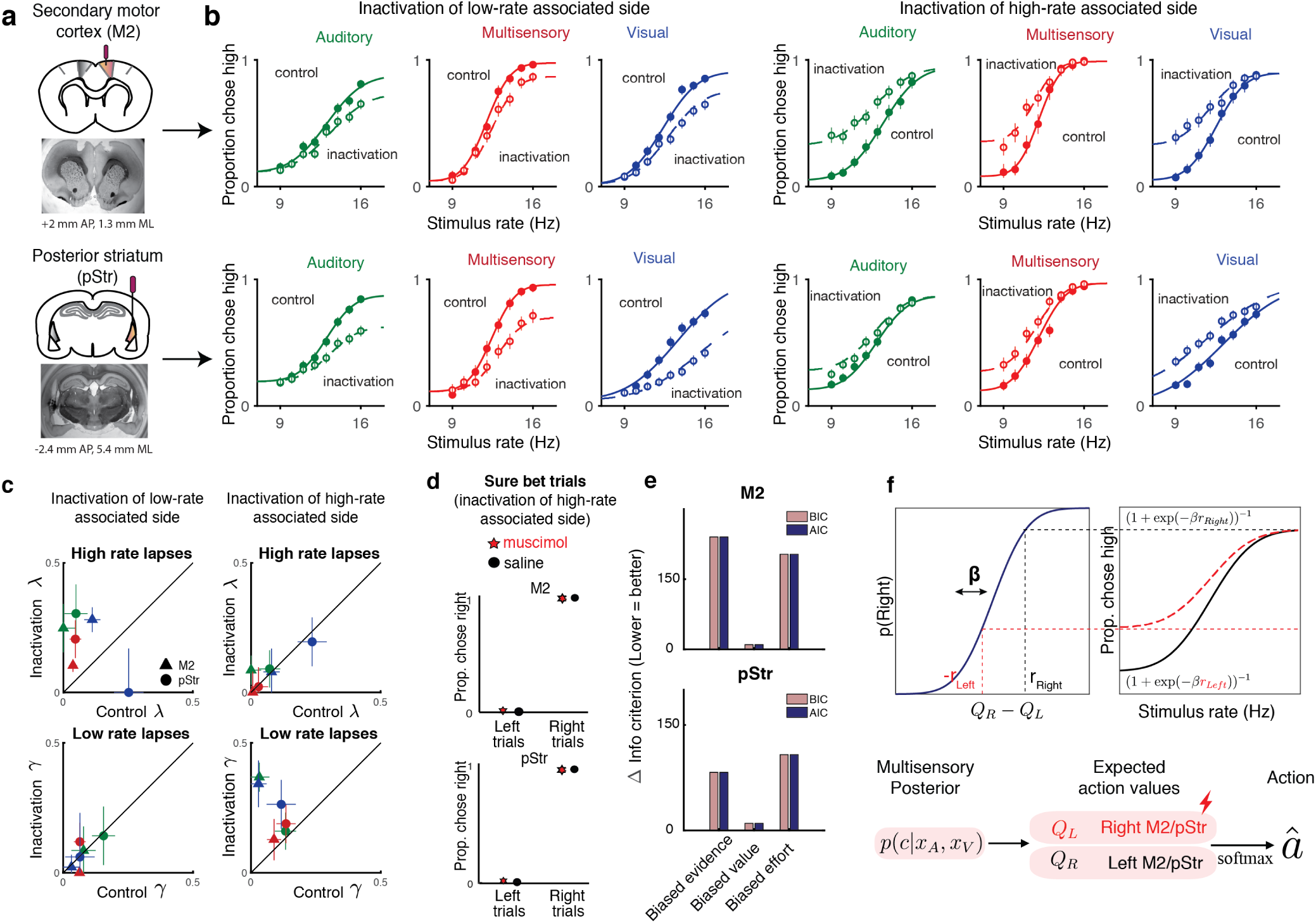
Inactivation of secondary motor cortex and posterior striatum affects lapses, suggesting a role in action value encoding. **(a)** Schematic of cannulae implants in M2 (top) and pStr (bottom) and representative coronal slices. For illustration purposes only, the schematic shows implants in the right hemisphere, however, the inactivations shown in panel (b) were performed unilaterally on both hemispheres. **(b)** Unilateral inactivation of M2 (top) and pStr (bottom). Left 6 plots: inactivation of the side associated with low-rates shows increased lapses for high rates on visual (blue), auditory (green) and multisensory (red) trials (M2: n=5 rats; 10329 control trials, full line; 6174 inactivation trials, dotted line; pStr: n=5 rats; 10419 control trials; 6079 inactivation trials). Right 6 plots: inactivation of the side associated with high-rates shows increased lapses for low rates on visual, auditory and multisensory trials (M2: n=3 rats; 5678 control trials; 3816 inactivation trials; pStr: n=6 rats; 11333 control trials; 6838 inactivation trials). Solid lines are exploration model fits, accounting for inactivation effects across all 3 modalities by scaling all contralateral values by a single parameter. **(c)** Increased high rate lapses following unilateral inactivation of the side associated with low-rates (top left); no change in low rate lapses (bottom left) and vice versa for inactivation of the side associated with high-rates (top, bottom right). Control data on the abscissa is plotted against inactivation data on the ordinate. Same animals as in **b**. Green, auditory trials; blue, visual trials; red, multisensory trials. Abbreviations: posterior striatum (pStr), secondary motor cortex (M2). **(d)** Sure bet trials are unaffected following inactivation. Pooled data shows that rats that were inactivated on the side associated with high rates make near perfect rightward and leftward choices Top, M2 (3 rats); bottom, pStr (6 rats). **(e)** Model comparison of three possible multisensory deficits - reduction of contralateral evidence by a fixed amount (left), reduction of contralateral value by a fixed amount (center) or an increased contralateral effort by a fixed amount (right). Both AIC and BIC suggest a value deficit **(f)** Proposed computational role of M2 and Striatum. Lateralized encoding of left and right action values by right and left M2/pStr (bottom) explains the asymmetric effect of unilateral inactivations on lapses (top).

Fitting the data with the exploration model revealed that, in keeping with the first model prediction, the effects on lapses in all modalities could be captured by scaling the contralateral action value by a single parameter (Fig. 5b, joint fits to control [solid lines] and inactivation trials [dotted lines] across modalities, differing only by a single parameter), similar to the reward manipulation experiments. Animals that were inactivated on the side associated with high rates showed increased lapses on low-rate trials (Fig. 5c, bottom right; data points are above the unity line; n=9 rats), but unchanged lapses on high-rate trials (Fig. 5c, top right; data points are on the unity line). This was consistent across areas and modalities (Fig. 5c; M2, triangles; pStr, circles; blue, visual; green, auditory; red, multisensory). Similarly, animals that were inactivated on the side associated with low rates showed the opposite effect: increased lapses on high-rate trials (Fig. 5c, top left; n=10 rats), while lapses did not change for low-rate trials (Fig. 5c bottom left). The effect was consistent across individual animals (M2, Supplementary Fig. 8; pStr, Supplementary Fig. 9).

In keeping with the second prediction, when we compared the effects of the disruptions in animals trained on standard and reversed contingencies (low rates rewarded with leftward or rightward actions respectively), the effects were always restricted to lapses on the stimuli associated with the side contralateral to the inactivation (Supplementary Fig. 10), always resembling a devaluation of contralateral actions (Supplementary Fig. 11).

A model comparison revealed that a fixed reduction in contralateral value captured the inactivation effects much better than a fixed reduction in contralateral sensory evidence or a fixed increase in contralateral motor effort, both for M2 (Fig. 5e top) and pStr (Fig. 5e bottom). In uncertain conditions, this reduced contralateral value gives rise to more exploratory choices and hence more lapses on one side (Fig. 5f top).

The final prediction of the exploration model is that changes in action value will only affect trials in which there was uncertainty about the outcome. In keeping with that prediction, performance was spared on sure-bet trials (Fig. 5d): rats made correct rightward and leftward choices regardless of the side that was inactivated. This observation provides further reassurance that the changes we observed on more uncertain conditions did not simply reflect motor impairments that drove a tendency to favor ipsilateral movements. Additional movement parameters such as wait time in the center port and movement times to ipsilateral and contralateral reward ports were likewise largely spared (Supplementary figure 12), suggesting that effects on decision outcome were not due to an inactivation-induced motor impairment.

Together, these results demonstrate that lapses are a powerful tool for interpreting behavioral changes in disruption experiments. For M2 and pStr disruptions, our analysis of lapses and deployment of the exploration model allowed us to reconcile previous inactivation studies. Our results suggest that M2 and pStr have a lateralized, modality-independent role in computing the expected value of actions (Fig. 5f bottom).

## DISCUSSION

Perceptual decision-makers have long been known to display a small fraction of errors even on easy trials. Until now, these “lapses” were largely regarded as a nuisance and lacked a comprehensive, normative explanation. Here, we propose a novel explanation for lapses: that they reflect a strategic balance between exploiting known rewarding options and exploring uncertain ones. Our model makes strong predictions for lapses under diverse decision-making contexts, which we have tested here. First, the model predicts more lapses on conditions with higher uncertainty, such as unisensory (Fig. 2) or neutral (Fig. 3), compared to multisensory or sure-bet conditions. Second, the model predicts that asymmetric reward manipulations should only affect lapses on one side, sparing decisions to the other side as well as sure-bet trials (Fig. 4). Finally, the model predicts that lapses should be affected by perturbations to brain regions that encode action value. Accordingly, we observed that unilateral inactivations of secondary motor cortex and posterior striatum similarly affected lapses on one side across auditory, visual and multisensory trials (Fig. 5). Taken together, our model and experimental data argue strongly that far from being a nuisance, lapses are informative about animals’ subjective action values and reflect a trade-off between exploration and exploitation.

Considerations of value have provided many useful insights into aspects of behavior that seem sub-optimal at first glance from the perspective of perceptual ideal observers. For instance, many perceptual tasks are designed with accuracy in mind - defining an ideal observer as one who maximizes accuracy, in line with classical signal detection theory. However, in practice, the success or failure of different actions may be of unequal value to subjects, especially if reward or punishment is delivered explicitly, as is often the case with non-human subjects. This may give rise to biases that can only be explained by an observer that maximizes expected utility (Dayan and Daw, 2008). Similarly, outcomes on a given trial can influence decisions about stimuli on subsequent trials through reinforcement learning, giving rise to serial biases. These biases occur even though the ideal observer should treat the evidence on successive trials as independent (Busse et al., 2011; Lak et al., 2018; Mendonca et al., 2018). When subjects can control how long they sample the stimulus, subjects maximizing reward rate may choose to make premature decisions, sacrificing accuracy for speed (Bogacz et al., 2006; Drugowitsch, DeAngelis, et al., 2014). Finally, additional costs of exercising mental effort could lead to bounded optimality through “satisficing” or finding good enough solutions (Mastrogiorgio and Petracca, 2018; Fan, Gold, and Ding, 2018).

Here, we take further inspiration from considerations of value to provide a novel, normative explanation for lapses in perceptual decisions. Our results argue that lapses are not simply accidental errors made as a consequence of attentional “blinks” or motor “slips”, but can reflect a deliberate, internal source of behavioral variability that facilitates learning and information gathering when the values of different actions are uncertain. This explanation connects a well known strategy in value-based decision making to a previously mysterious phenomenon in perceptual decision making.

Although exploration no longer yields the maximum utility on any given trial, it is critical for dynamic environments, and those in which there is uncertainty about probability of reward or stimulus-response contingency, especially if these need to be learnt or refined through experience. By encouraging subjects to sample multiple options, exploration can potentially improve subjects’ knowledge of the rules of the task, helping them to increase future payoff, thus maximizing expected utility over a long period of time. This offers an explanation for the higher rate of lapses seen in humans on tasks with abstract (Raposo, Sheppard, et al., 2012), non-intuitive (Mihali et al., 2018) or non-verbalizable (Flesch et al., 2018) stimulus-response contingencies.

Balancing exploration and exploitation is computationally challenging, and the mechanism we propose here, Thompson sampling, is an elegant heuristic for achieving this balance. This strategy has been shown to be asymptotically optimal in partially observable environments (Leike et al., 2016) and can be naturally implemented through a sampling scheme where the subject samples action values from a learnt distribution and then maximizes with respect to the sample (Gershman, 2018). This strategy predicts that conditions with higher uncertainty should have higher exploration, and consequently higher lapse rates, explaining the pattern of lapse rates we observed on unisensory vs. multisensory trials as well as on neutral vs. matched trials. A lower rate of lapses on multisensory trials has also been reported on a visual-tactile task in rats (Nikbakht et al., 2018) and a vestibular integration task in humans (Bertolini et al., 2015) and can potentially account for the apparent supra-optimal integration that has been reported in a number of rodent, non-human primate and human studies (Nikbakht et al., 2018; Hou et al., 2018; Raposo, Sheppard, et al., 2012). A strong prediction of uncertainty guided exploration is that the animal should quickly learn to exploit on conditions with no uncertainty, as we observed on sure-bet trials (Fig. 4d, 5d).

Uncertainty-guided exploration also predicts that exploratory choices, and consequently lapses, should decrease with training as the animal becomes more certain of the rules and expected rewards, explaining training-dependent effects reported in primates (Law and Gold, 2009; Cloherty et al., 2019). This can also potentially explain why children have higher lapse rates (Witton, Talcott, and Henning, 2017; Manning et al., 2018)- they have been shown to be more exploratory in their decisions than adults (Lucas et al., 2014).

A unique prediction of the exploration model is that it predicts lapse rates will sometimes change asymmetrically for left and right decisions. For instance, changing the value associated with one of the decisions (eg. high rate) should only affect lapses associated with that decision - predicting more lapses on high rates if the rightward reward is decreased or if leftward decisions are probabilistically rewarded on high rates. Similarly, increasing rightward rewards should decrease lapses on high rates. These predictions are borne out (Fig. 4c), and rightward successes or failures on the previous trial have a similar effect. The model also suggests that the asymmetric effects on lapses seen during unilateral inactivations of prefrontal and striatal regions (Fig. 5b) arise from a selective devaluation of contralateral actions. This interpretation reconciles a number of studies that have found asymmetric effects of inactivating these areas during perceptual decisions (Erlich et al., 2015; Zatka-Haas et al., 2019; Wang et al., 2018; Guo et al., 2018) with their established roles in encoding action value (Barthas and Kwan, 2017; Lee et al., 2015) during value-based decisions.

An open question that remains is how the brain might tune the degree of exploration in proportion to uncertainty. An intriguing candidate for this is dopamine, whose phasic responses have been shown to reflect state uncertainty (Starkweather et al., 2017; Babayan, Uchida, and Gershman, 2018; Lak et al., 2018), and whose tonic levels have been shown to modulate exploration in mice on a lever-press task (Beeler et al., 2010), and context-dependent song variability in songbirds (Leblois, Wendel, and Perkel, 2010). Dopaminergic genes have been shown to predict individual differences in uncertainty-guided exploration in humans (Frank et al., 2009), and dopaminergic disorders such as Parkinson’s disease have been shown to disrupt the uncertainty-dependence of lapses across conditions on a multisensory task (Bertolini et al., 2015). Patients with ADHD, another disorder associated with dopaminergic dysfunction, have been shown to display both increased perceptual variability and increased task-irrelevant motor output, a measure that correlates with lapses (Mihali et al., 2018). A promising avenue for future studies is to leverage the informativeness of lapses and the precise control of uncertainty afforded by multisensory tasks, in conjunction with perturbations or recordings of dopaminergic circuitry, to further elucidate the connections between perceptual and value-based decision making systems.

## METHODS

### Behavior

#### Animal Subjects and Housing

All animal procedures and experiments were in accordance with the National Institutes of Healths Guide for the Care and Use of Laboratory Animals and were approved by the Cold Spring Harbor Laboratory Animal Care and Use Committee. Experiments were conducted with 34 adult male and female Long Evans rats (250-350g, Taconic Farms) that were housed with free access to food and restricted access to water starting from the onset of behavioral training. Rats were housed on a reversed light-dark cycle; experiments were run during the dark part of the cycle. Rats were pair-housed during the whole training period.

#### Animal training and behavioral task

Rats were trained following previously established methods (Raposo 2012, Sheppard 2013, Raposo 2014, Licata 2017). Briefly, rats were trained to wait in the center port for 1000 ms while stimuli were presented, and to associate stimuli with left/right reward ports. Stimuli for each trial consisted of a series of events: auditory clicks from a centrally positioned speaker, full-field visual flashes, or both together. Stimulus events were separated by either long (100 ms) or short (50 ms) intervals. For the easiest trials, all inter-event intervals were identical, generating rates that were 9 events/s (all long intervals) or 16 events/s (all short intervals). More difficult trials included a mixture of long and short intervals, generating stimulus rates that were intermediate between the two extremes and therefore more difficult for the animal to judge. The stimulus began after a variable delay following when the rats snout broke the infrared beam in the center port. The length of this delay was selected from a truncated exponential distribution (*λ* = 30 ms, minimum = 10 ms, maximum = 200 ms) to generate an approximately flat hazard function. The total time of the stimulus was usually 1000 ms. Trials of all modalities and stimulus strengths were interleaved. For multisensory trials, the same number of auditory and visual events were presented (except for a subset of neutral trials). Auditory and visual stimulus event times were generated independently, as our previous work has demonstrated that rats make nearly identical decisions regardless of whether stimulus events are presented synchronously or independently (Raposo, Sheppard, et al., 2012). For most experiments, rats were rewarded with a drop of water for moving to the left reward port following low-rate trials and to the right reward port following high rate trials. For muscimol inactivation experiments half of the rats were rewarded according to the reverse contingency. Animals typically completed between 700 and 1,200 trials per day. Most experiments had 18 conditions (3 modalities, 8 stimulus strengths), leading to 29-50 trials per condition per day.

To probe the effect of uncertainty on lapses, rats received catch trials consisting of multisensory neutral trials, where only the auditory modality provided evidence for a particular choice, whereas the visual modality provided evidence that was so close to the category boundary (12 Hz) that it did not support one choice or the other (Raposo, Sheppard, et al., 2012)

To probe the effect of value on lapses, we manipulated either reward magnitude or reward probability associated with high rates, while keeping low rate trials unchanged. To increase or decrease reward magnitude associated with high rates, the amount of water dispensed on the right port was increased or decreased to 36*µ*l or 16*µ*l respectively, while the reward on the left port was maintained at 24 *µ*l. To manipulate reward probability, we occasionally rewarded rats on the (incorrect) left port on high rate trials with a probability of 0.5. The right port was still rewarded with a probability of 1 on high rates, and reward probabilities on low rate trials were unchanged (1 on the left port, 0 on the right).

### Analysis of behavioral data

#### Psychometric curves

Descriptive four-parameter psychometric functions were fit to choice data using the Palamedes toolbox (Prins and Kingdom, 2018). Psychometric functions were parameterized as:

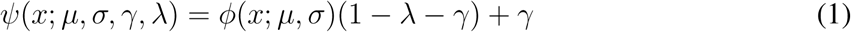

where *γ* and *λ* are the lower and upper asymptote of the psychometric function, which parametrize the lapse rates on low and high rates, respectively. *ϕ* is a cumulative normal function; *x* is the event rate, i.e. the number of flashes or beeps presented during the one second stimulus period; *µ* parametrizes the x-value at the midpoint of the psychometric function and *σ* describes the inverse slope. 95% Confidence intervals on these parameters were generated via bootstrapping based on 1000 simulations.

Our definition of lapses is restricted to strictly *asymptotic* errors following Wichmann and Hill, 2001, and not simply errors on the easiest stimuli tested. Errors on the easiest stimuli could in general arise not just from lapses (strictly defined) but also from perceptual errors caused by low sensitivity to the stimulus, an insufficient stimulus range or non-stationary weights (Busse et al., 2011; Roy et al., 2018). However we do not consider easy errors alone to be evidence of lapses and only consider asymptotic errors. To confirm the necessity of including the lapse parameters, we fit variants of the model above where the lapse parameters are restricted to be zero, and only included them when warranted by model comparison using AIC/BIC.

### Modeling

#### Ideal observer model

We can specify an ideal observer model for our task using Bayesian Decision Theory (Dayan and Daw, 2008). This observer maintains probability distributions over previously experienced stimuli and choices, computes the posterior probability of each action being correct given its observations and picks the action that yields the highest expected reward.

Let the true category on any given trial be *c*_*true*_, the true stimulus rate be *s*_*true*_ and the animal’s noisy visual and auditory observations of *s*_*true*_ be *x*_*V*_ and *x*_*A*_, respectively. We assume that the two sensory channels are corrupted by independent gaussian noise with standard deviation *σ*_*A*_ and *σ*_*V*_, respectively, giving rise to conditionally independent observations.

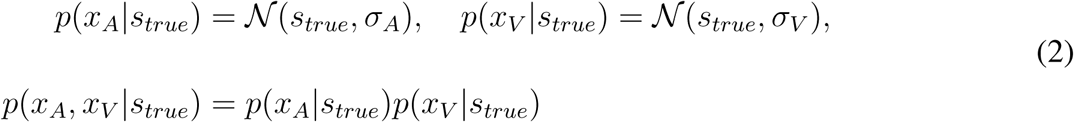

The ideal observer can use this knowledge to compute the likelihood of seeing the current trial’s observations as a function of the hypothesized stimulus rate *s*. This likelihood *ℒ* is a gaussian function of *s* with a mean given by a weighted sum of the observations *x*_*A*_ and *x*_*V*_,:

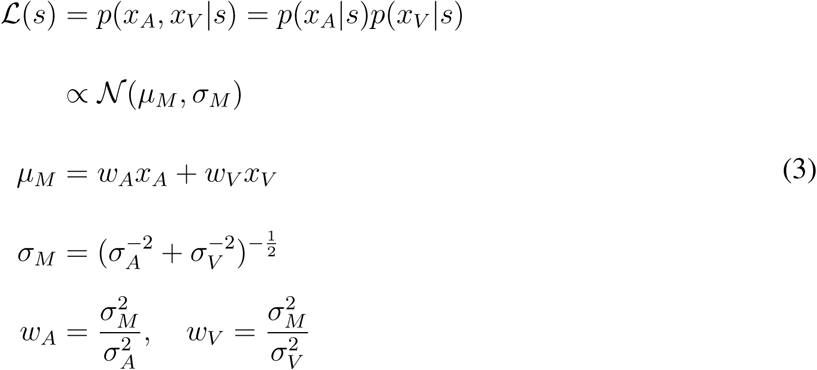

The likelihood of seeing the observations as a function of the hypothesized category *c*, is given by marginalizing over all possible hypothesized stimulus rates. Let the experimentally imposed category boundary be *µ*_0_, such that stimulus rates are considered high when *s* > *µ*_0_ and low when *s* < *µ*_0_. Then,

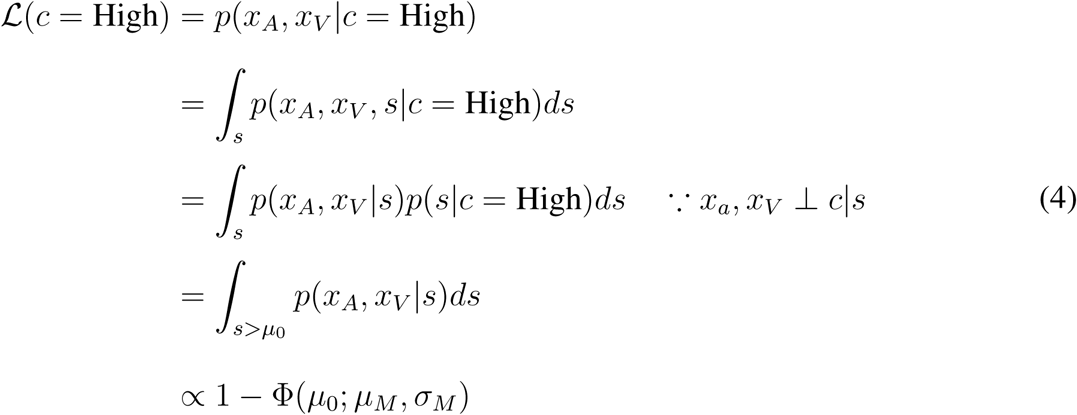

where Φ is the cumulative normal function. Using Bayes’ rule, the ideal observer can then compute the probability that the current trial was high or low rate given the observations, i.e. the posterior probability.

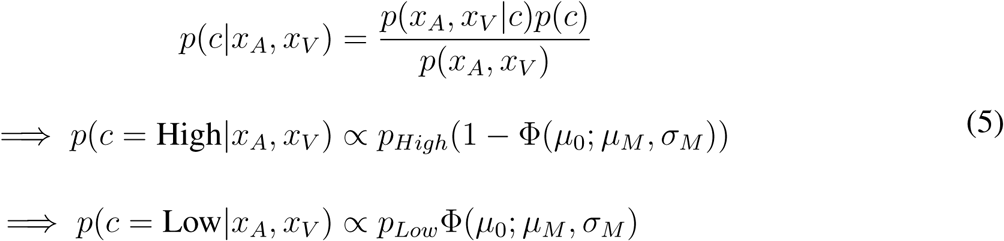

where *p*_*High*_ and *p*_*Low*_ are the prior probabilities of high and low rates respectively. The expected value *Q*(*a*) of choosing right or left actions (also known as the action values) is obtained by marginalizing the learnt value of state-action pairs *q*(*c, a*) over the unobserved state *c*.

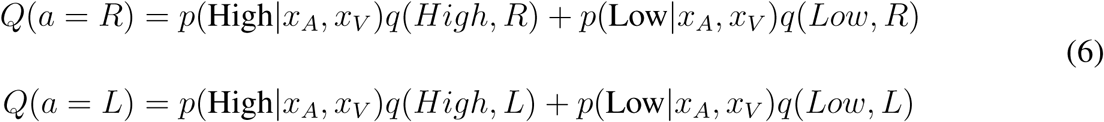

Under the standard contingency, high rates are rewarded on the right and low rates on the left, so for a trained observer that has fully learnt the contingency, *q*(*High, R*) → *r*_*R*_, *q*(*High, L*) → 0, *q*(*Low, R*) → 0, *q*(*Low, L*) → *r*_*L*_, with *rR* and *rL* being reward magnitudes for rightward and leftward actions. This simplifies the action values to:

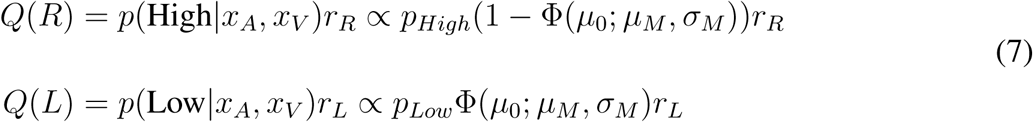

The max-reward decision rule involves picking the action *â* with the highest expected reward:

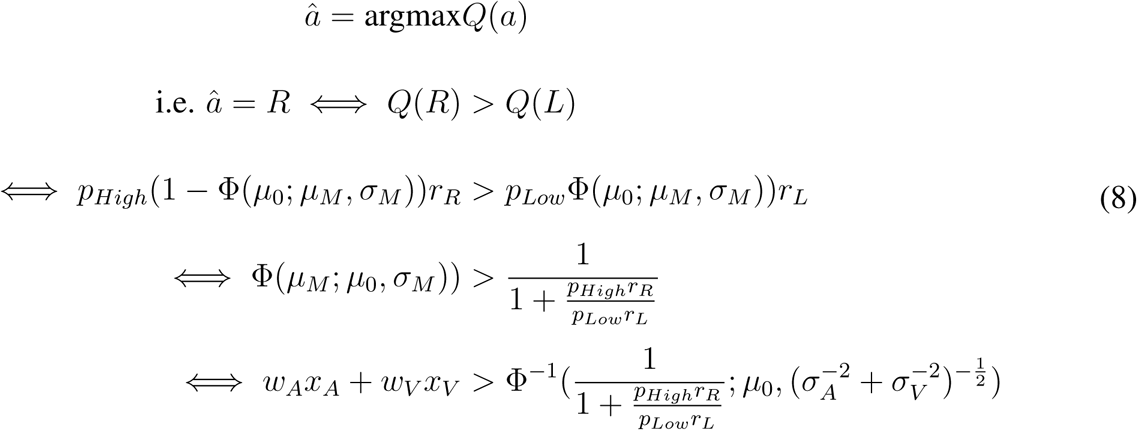

In the special case of equal rewards and uniform stimulus and category priors, this reduces to choosing right when the weighted sum of observations is to the right of the true category boundary, i.e. *w*_*A*_*x*_*A*_ + *w*_*V*_ *x*_*V*_ > *µ*_0_. Note that this is a deterministic decision rule for any given observations *x*_*A*_ and *x*_*V*_, however, since these are noisy and gaussian distributed around the true stimulus rate *s*_*true*_, the likelihood of making a rightward decision is given by the cumulative gaussian function Φ:

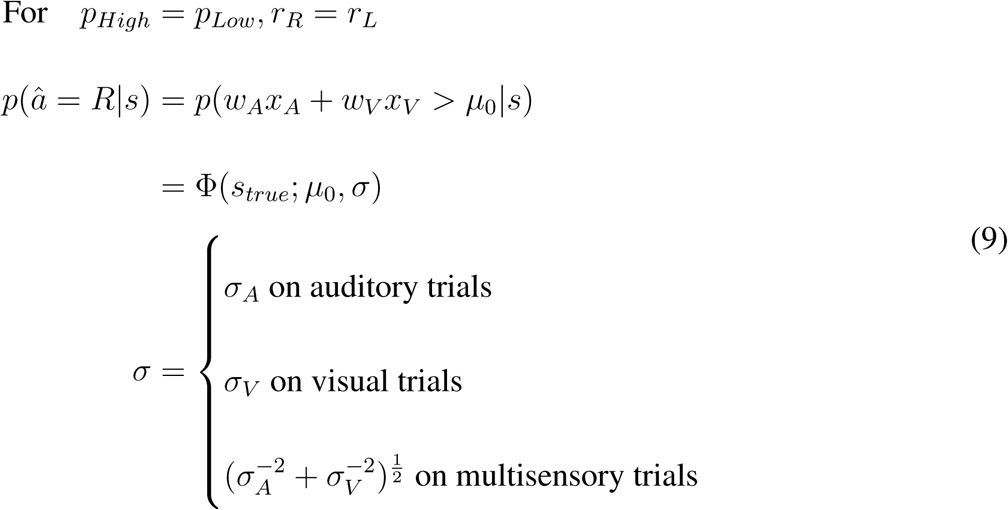

We can measure this probability empirically through the psychometric curve. Fitting it with a two parameter cumulative gaussian function yields *µ* and *σ* which can be compared to ideal observer predictions. The *σ* parameter is then taken to reflect sensory noise; and with the assumption of uniform priors and equal rewards, the *µ* parameter is taken to reflect the subjective category boundary. Although *µ* should equal *µ*_0_ for the ideal observer, in practice it is treated as a free parameter, and deviations of *µ* from *µ*_0_ could reflect any of three possible suboptimalities: 1) a subjective category boundary mismatched to the true one, 2) mismatched priors, or 3) unequal subjective rewards of the two actions.

#### Inattention model

The traditional model for lapse rates assumes that on a fixed proportion of trials, the animal fails to pay attention to the stimulus, guessing randomly between the two actions. We can incorporate this suboptimality into the ideal observer above as follows: Let the probability of attending be *p*_*attend*_. Then, on 1 − *p*_*attend*_ fraction of trials, the animal does not attend to the stimulus (i.e. receives no evidence), effectively making *σ*_*sensory*_ → *∞* and giving rise to a posterior that is equal to the prior. On these trials, the animal may choose to maximize this prior (always picking the option that’s more likely a-priori, guessing with 50-50 probability if both options are equally likely), or probability-match the prior (guessing in proportion to its prior). Let us call this guessing probability *p*_*bias*_. Then, the probability of a rightward decision is given by marginalizing over the attentional state:

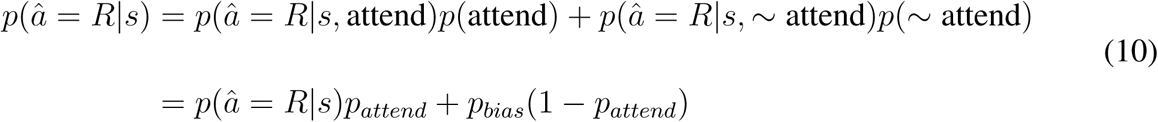

Comparing this with the standard 4-parameter sigmoid used in psychometric fitting, we obtain

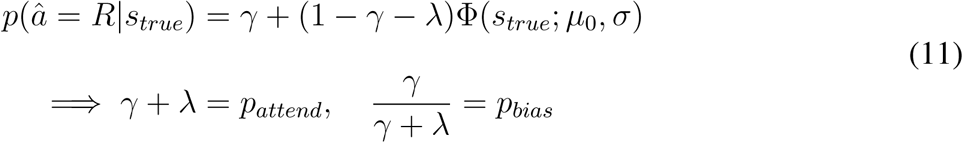

where *γ* and *λ* are the lower and upper asymptotes respectively, collectively known as “lapses”. In this model, the sum of the two lapses depends on the probability of attending, which could be modulated in a bottom up fashion by the salience of the stimulus; their ratio depends on the guessing probability, which in turn depends on the observer’s priors and subjective rewards.

#### Motor error/ϵ greedy model

Lapses can also occur if the observer doesn’t always pick the reward-maximizing or “exploit” decision. This might occur due to random errors in motor execution on a small fraction of trials given by *ϵ*, or it might reflect a deliberate propensity to occasionally make random “exploratory” choices to gather information about rules and rewards. This is known as an *ϵ*-greedy decision rule, where the observer chooses randomly (or according to *p*_*bias*_) on *ϵ* fraction of trials. Both these models yield predictions similar to those of the inattention model:

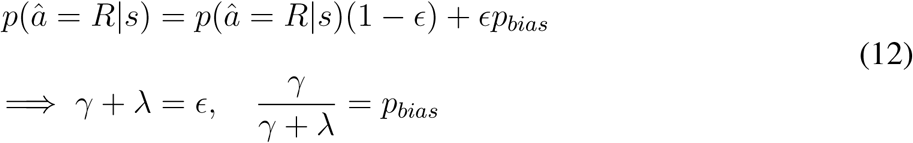

#### Uncertainty guided exploration model

A more sophisticated form of exploration is the “softmax” decision rule, which explores options in proportion to their expected rewards, allowing for a balance between exploration and exploitation through the tuning of a parameter *β* known as inverse temperature. In particular, in conditions of greater uncertainty about rules or rewards, it is advantageous to be more exploratory and have a lower *β*. This form of uncertainty-guided exploration is known as Thompson sampling. It can be implemented by sampling from a belief distribution over expected rewards and maximizing with respect to the sample, reducing to a softmax rule whose *β* depends on the total uncertainty in expected reward (Gershman, 2018).

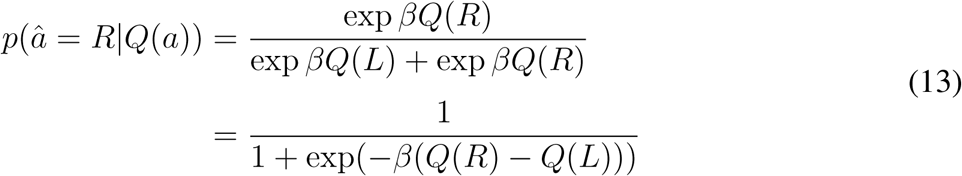

The proportion of rightward choices conditioned on the true stimulus rate is then obtained by marginalizing over the latent action values *Q*(*a*), using the fact that the choice depends on *s* only through its effect on *Q*(*a*), where *ρ* is the animal’s posterior belief in a high rate stimulus, i.e. *ρ* = *p*(*c* = *High*|*x*_*A*_, *x*_*V*_). *ρ* is often referred to as the *belief state* in reinforcement learning problems involving partial observability such as our task.

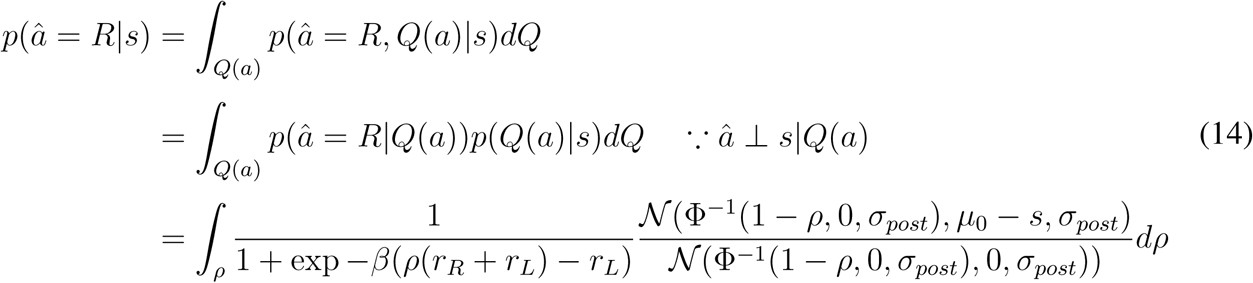

Since lapses are the asymptotic probabilities of the lesser rewarding action at extremely easy stimulus rates, we can derive them from this expression by setting *ρ* → 1 or *ρ* → 0. This yields

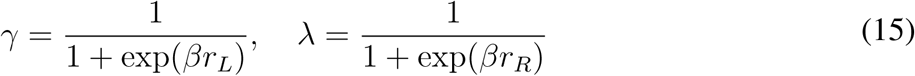

Critically, in this model, the upper and lower lapses are dissociable, depending only on the rightward or leftward rewards, respectively. Such a softmax decision rule has been used to account for suboptimalities in value based decisions (Dayan and Daw, 2008), however it has not been used to account for lapses in perceptual decisions. Other suboptimal decision rules described in perceptual decisions, such as generalized probability matching or posterior sampling (Acerbi, Vijayakumar, and Wolpert, 2014; Drugowitsch, Wyart, et al., 2016; Ortega and Braun, 2013) amount to a softmax on log-posteriors or log-expected values, rather than on expected values, and do not produce lapses since in these decision rules, when the posterior probability goes to 1, so does the decision probability.

### Model fitting

Model fits were obtained from custom maximum likelihood fitting code using MATLAB’s fmincon, by maximizing the marginal likelihood of rightward choices given the stimulus on each trial as computed from each model. Confidence intervals for fit parameters were generated using the hessian obtained from fmincon. Fits to multiple conditions were performed jointly, taking into account any linear or nonlinear (eg. optimality) constraints on parameters across conditions. Model comparisons were done using AIC and BIC.

### Surgical procedures

All rats subject to surgery were anesthetized with 1%-3% isoflurane. Isoflurane anesthesia was maintained by monitoring respiration, heart rate, oxygen and CO_2_ levels, as well as foot pinch responses throughout the surgical procedure. Ophthalmic ointment was applied to keep the eyes moistened throughout surgery. After scalp shaving, the skin was cleaned with 70% ethanol and 5% betadine solution. Lidocaine solution was injected below the scalp to provide local analgesia prior to performing scalp incisions. Meloxicam (5mg/ml) was administered subcutaneously (2mg/kg) for analgesia at the beginning of the surgery, and daily 2-3 days post-surgery. The animals were allowed at least 7 days to recover before behavioral training.

#### Viral injections

2 rats, 15 weeks of age, were anesthetized and placed in a stereotaxic apparatus (Kopf Instruments). Small craniotomies were made in the center of primary visual cortex (V1; 6.9mm posterior to Bregma, 4.2mm to the right of midline) and primary auditory cortex (A1; 4.7mm posterior to Bregma, 7mm to the right of midline). Small durotomies were performed at each craniotomy and virus was pressure injected at depths of 600, 800, and 1000 *µ*m below the pia (150 nL/depth). Virus injections were performed using Drummond Nanoject III, which enables automated delivery of small volumes of virus. To minimize virus spread, the Nanoject was programmed to inject slowly: fifteen 10 nL boluses, 30 seconds apart. Each bolus was delivered at 10 nL/sec. 2-3 minutes were allowed following injection at each depth to allow for diffusion of virus. The AAV2.CB7.CI.EGFP.WPRE.RBG construct was injected in V1, and the AAV2.CAG.tdTomato.WPRE.SV40 construct was injected in A1. Viruses were obtained from the University of Pennsylvania vector core.

#### Cannulae implants

Rats were anesthetized and placed in the stereotax as described above. After incision and skull cleaning, 2 skull screws were implanted to add more surface area for the dental cement. For striatal implants, two craniotomies were made, one each side of the skull (3.2mm posterior to Bregma; 5.4mm to the right and left of midline). Durotomies were performed and a guide cannula (22 gauge, 8.5 mm long; PlasticsOne) was placed in the brain, 4.1mm below the pia at each craniotomy. For secondary motor cortex implants, one large craniotomy spanning the right and left M2 was performed (∼5mm × ∼2mm in size centered around 2mm anterior to Bregma and 3.1mm to the right and left of midline). A durotomy was performed and a double guide cannula (22 gauge, 4mm long; PlasticsOne) was placed in the brain, 300*µ*m below the pia. The exposed brain was covered with sterile Vaseline and cannulae were anchored to the skull with dental acrylic (Relyx). Single or double dummy cannulae protruding 0.7 mm below the guide cannulae were inserted.

### Inactivation with muscimol

Rats were lightly anesthetized with isoflurane. Muscimol was unilaterally infused into pStr or M2 with a final concentration of 0.075-0.125 *µ*g and 0.1-0.5 *µ*g, respectively. A single/double-internal cannula (PlasticsOne), connected to a 2 *µ*l syringe (Hamilton microliter syringe, 7000 series), was inserted into each previously implanted guide cannula. Internal cannulae protruded 0.5mm below the guide. Muscimol was delivered using an infusion pump (Harvard PHD 22/2000) at a rate of 0.1 *µ*l/minute. Internal cannulae were kept in the brain for 3 additional minutes to allow for diffusion of muscimol. Rats were removed from anesthesia and returned to cages for 15 minutes before beginning behavioral sessions. The same procedure was used in control sessions, where muscimol was replaced with sterile saline.

### Histology

At the conclusion of inactivation experiments, animals were deeply anesthetized with Euthasol (pentobarbital and phenytoin). Animals were perfused transcardially with 4% paraformaldehyde. Brains were extracted and post-fixed in 4% paraformaldehyde for 24-48 hours. After post-fixing, 50-100 *µ*m coronal sections were cut on a vibratome (Leica) and imaged.

## Acknowledgements

We thank Matt Kaufman, Simon Musall, Onyekachi Odoemene, Ashley Juavinett, Farzaneh Najafi, Akihiro Funamizu, Priyanka Gupta, Anne Urai, James Roach, Colin Stoneking, Diksha Gupta, Tatiana Engel, Rob Phillips, Tony Zador, Steve Shea and Bo Li for scientific advice and discussions, and Angela Licata, Steven Gluf, Liete Einchorn, Dennis Maharjan, Alexa Pagliaro, Edward Lu and Barry Burbach for technical assistance. We thank Partha Mitra, Alexander Tolpygo and Stephen Savoia for help with slicing and imaging virus injected brains. This work was supported by the Simons Collaboration on the Global Brain, ONR MURI, the Eleanor Schwartz Fund, the Pew Charitable Trust and the Watson School of Biological Sciences.

## Supplemental Text and Figures

**Supplementary Figure 1:**
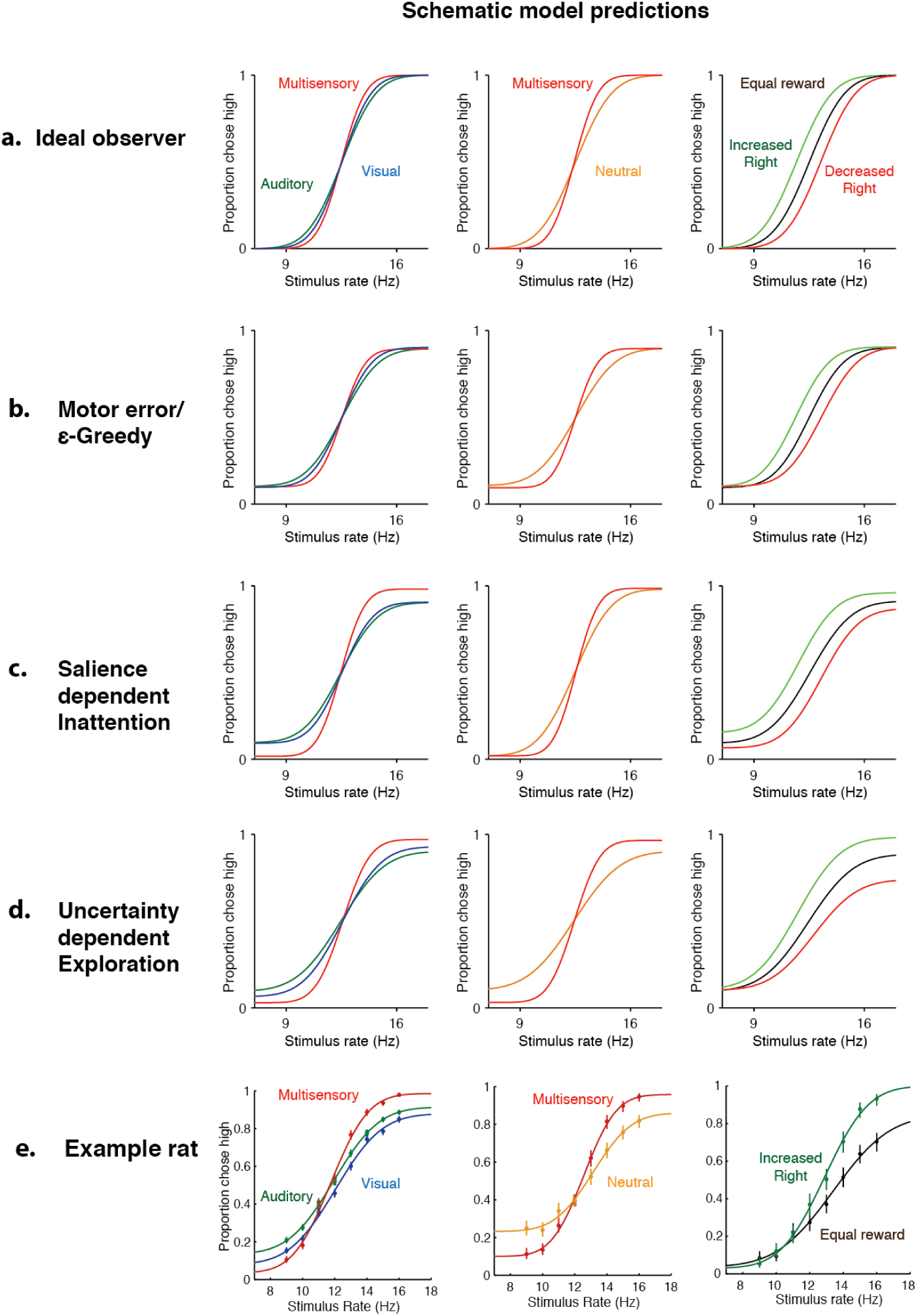
Uncertainty-dependent exploration is the only model that accounts for behavioral data from all three manipulations. Columns: data/predictions for three experimental manipulations. Left: unisensory vs. multisensory. Middle: matched vs. neutral. Right: Asymmetric reward. a-d: Four candidate models. (a) Ideal observer model predicts no lapses and only changes in sensitivity/bias across conditions. (b) Fixed motor error model predicts a constant rate of lapses across conditions in addition to changes in sensitivity/bias predicted from the ideal observer. (c) Inattention model predicts that the overall lapse rate (sum of lapses on both sides) depends on the level of bottom-up attentional salience, allowing for different rates for unisensory and multisensory. It also predicts that the lapse rate on neutral trials should be equal to that on multisensory trials, and that manipulating right-ward reward should affect both lapse rates. (d) Uncertainty-dependent exploration model predicts that overall lapse rate depends on the level of exploratoriness and hence uncertainty associated with that condition, allowing for different lapse rates on unisensory and multisensory trials. It also predicts that the lapse rate on neutral trials should be equal to that on auditory trials and manipulating rightward reward should only affect high rate lapses. (e) Data from an example rat on all three manipulations.

**Supplementary Figure 2:**
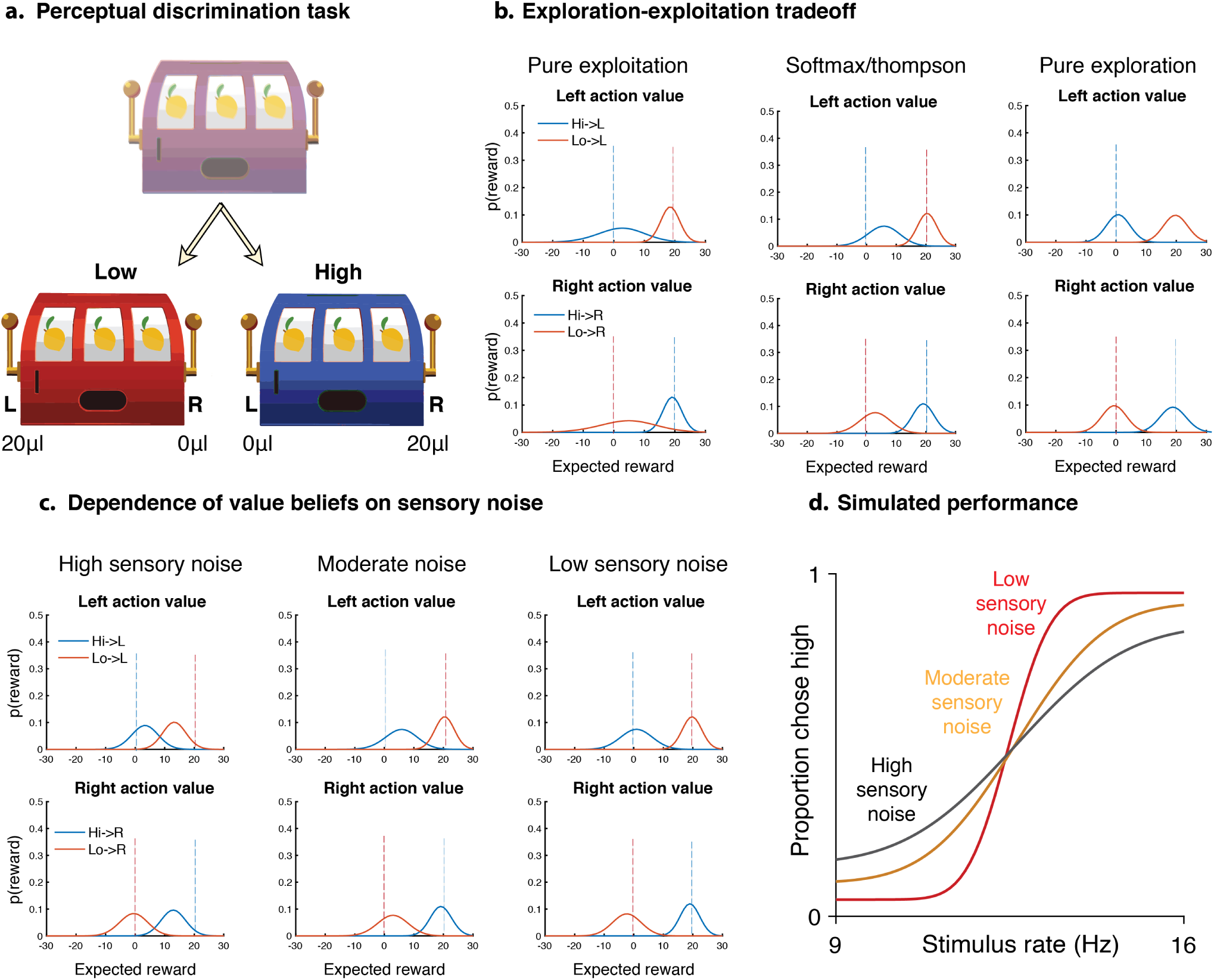
Thompson sampling, which balances exploration and exploitation, predicts lapses that increase with perceptual noise. Schematic illustrating the explore-exploit tradeoff in perceptual two-alternative tasks. (a) Formulation of perceptual decision making task as a partially observable contextual bandit. To solve this task, an observer needs to infer the true category of the stimulus (Low or High) based on noisy observations, and pick the best action given the true category (Left for Low, Right for High). This requires accurately learning the expected rewards from all 4 state-action pairs. (b) Beliefs about expected reward from different actions (L,R) performed in different states (Hi, Lo) showing different levels of uncertainty depending on policy. Beliefs are updated based on outcomes using a Bayesian update rule that takes into account uncertainty in state estimation. A greedy policy (left) that always picks the best action maximizes reward and learns well about the preferred state-action pairs (i.e. Lo-L and Hi-R) but has high uncertainty about the non-preferred pairs (Lo-R, Hi-L). A random policy (right) earns reward at chance, but learns equally well about all state-action pairs. Thompson sampling (center) implements a softmax decision rule that depends on the current uncertainties in each value, and balances immediately reward-maximizing decisions with decisions that reduce uncertainty, maximizing average reward in the long term. (c) Learnt beliefs about expected reward with Thompson sampling at various levels of perceptual uncertainty. High sensory noise (left) leads to large perceptual uncertainty, yielding highly overlapping belief distributions owing to a reduced ability to assign obtained rewards to one of the states. Lower levels of sensory noise (center, right) produce more separable beliefs. (d) Simulated performance over 2000 trials of the Bayesian learner shown above, under a Thompson sampling policy. As the sensory noise decreases (Black to Yellow to Red), the observer makes fewer exploratory choices owing to the more separable value beliefs, giving rise to lower lapse rates.

**Supplementary Figure 3:**
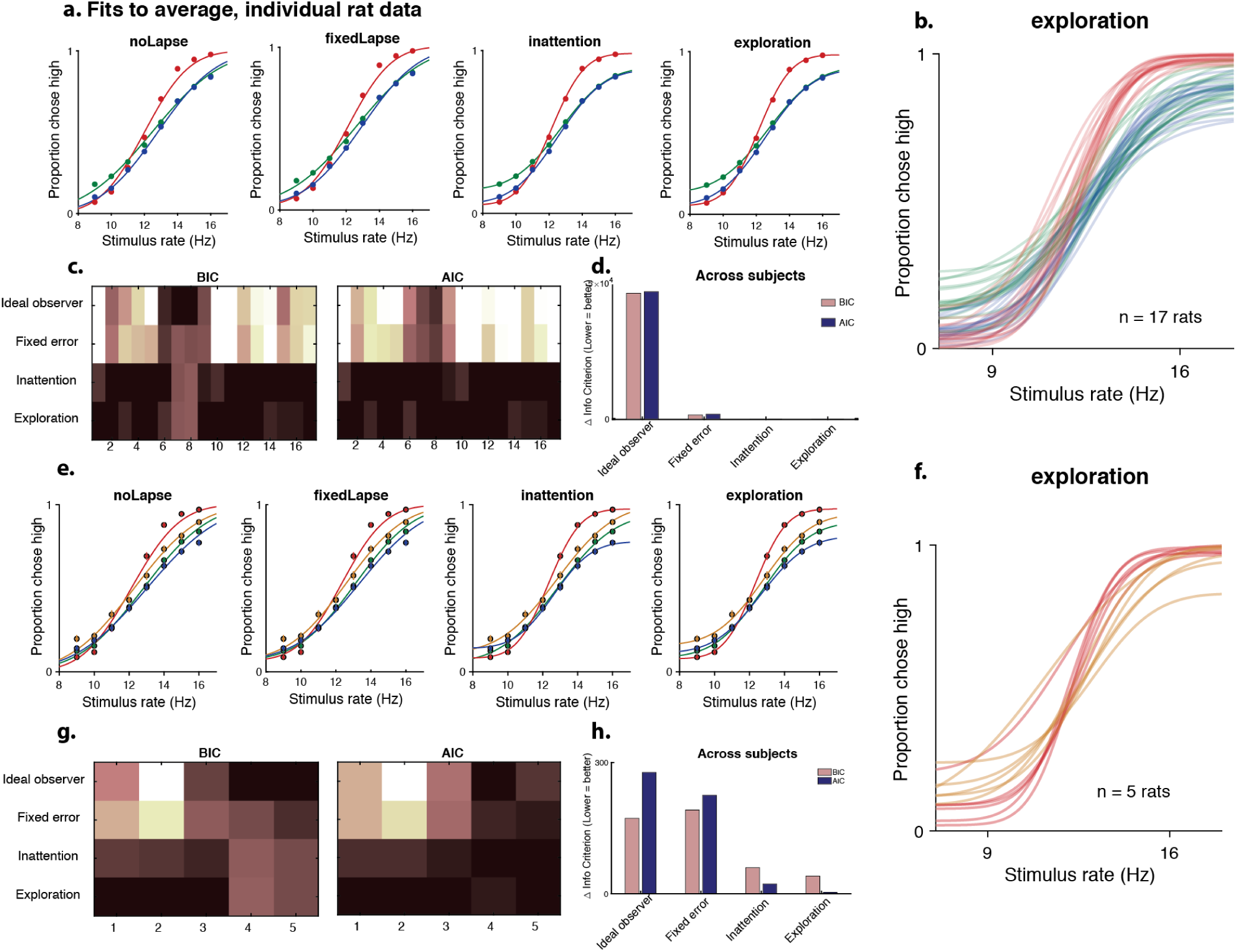
Uncertainty guided exploration outperforms competing models for average and individual data. (a) Fits of the four models to average rat data on unisensory (blue-visual, green-auditory) and multisensory (red) trials. (b) Exploration model fits to unisensory and multisensory data for 17 individual animals (c) Model comparison for individual animals using BIC (left), AIC (right). Darker colors are lower BICs/AICs, denoting a better fit. (d) Summed model comparison metrics across animals, showing that inattention and exploration models fit the data equally well, and much better than the ideal observer or fixed error models. (e) Fits of the four models to average data including neutral trials (orange) provide a stronger test of the inattention model. (f) Exploration model fits to multisensory data including neutral trials for 5 individual animals (g) Model comparison for individual animals. (h) Summed model comparison metrics across animals shows that the uncertainty-guided exploration model performs better than other models.

**Supplementary Figure 4:**
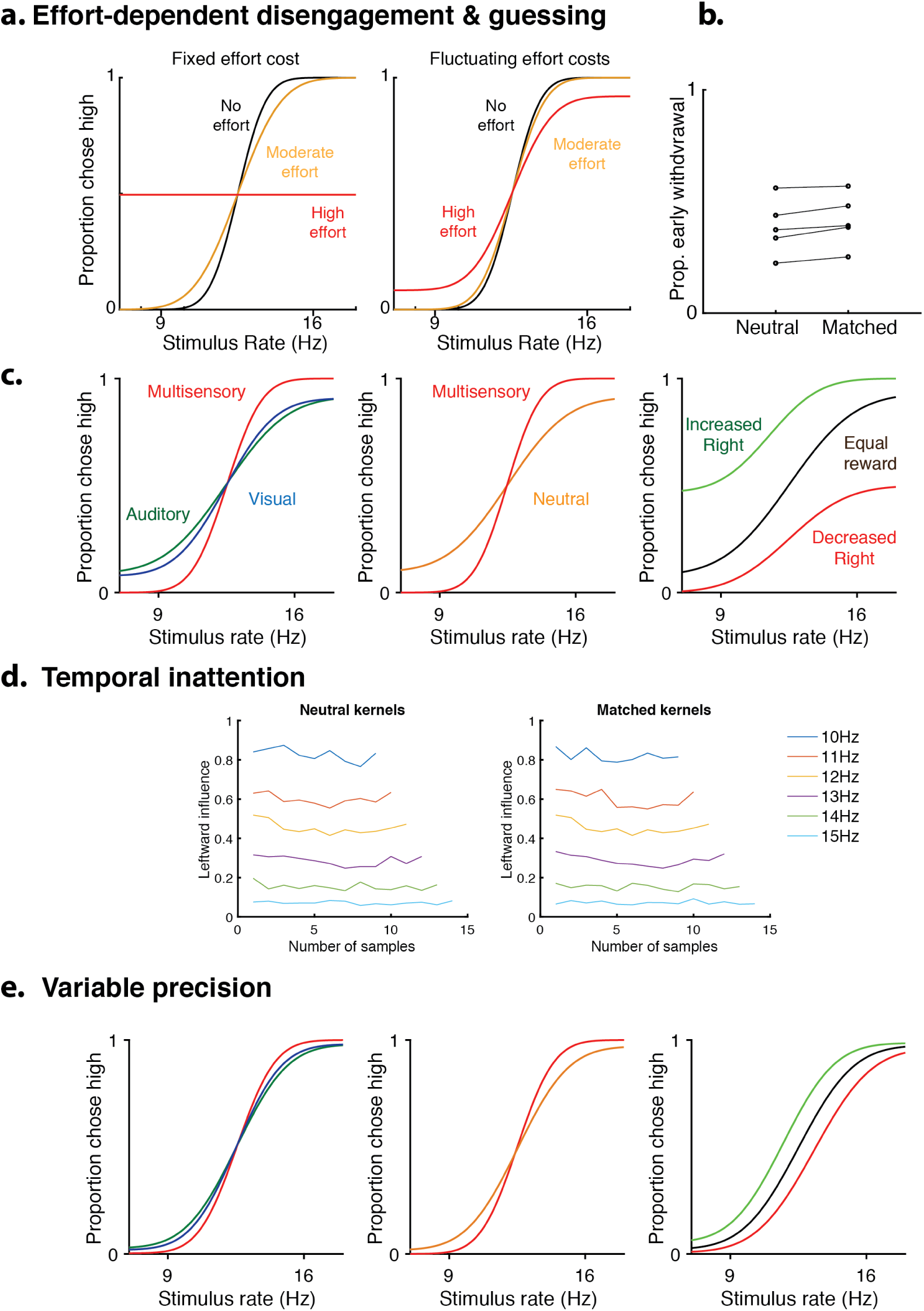
Alternative models of inattentional lapses. Predictions of alternative models of lapses. (a) Effort-dependent disengagement model: In this model, there is an additional cost or mental effort to being engaged in the task which could vary with condition, and an additional random guessing action. If the net payoff of engagement is not greater than the average value of a guess, then it guesses randomly. Such a model does not produce lapses if the effort is fixed across trials (left), but could produce lapses if the effort fluctuates from trial to trial (center). (b) Proportion of trials on which the animal withdrew prematurely doesnt vary between matched and neutral trials, suggesting that rats are not disengaging preferentially on neutral trials. (c) Predictions of the effort-dependent disengagement model. The model accurately predicts increased lapses on unisensory trials (left panel, green/blue traces) and neutral multisensory trials (middle panel, orange trace). However, for asymmetric reward manipulations (right), the model fails to predict our behavioral observation (Fig. 4d) that only lapses on the manipulated side are affected. (d) Temporal inattention model: in this model, temporal weighting of evidence differs between matched and neutral trials. To test this, we compared psychophysical kernels on matched and neutral trials. The temporal dynamics of attention are unchanged between the two kinds of trials, arguing against the temporal inattention model. (e) Variable precision model: in this model, the sensory noise (or its inverse, precision) fluctuates from trial to trial. The model accurately predicts increased lapses on unisensory trials (left panel, green/blue traces) and neutral multisensory trials (middle panel, orange trace). However, for asymmetric reward manipulations (right), the model fails to predict our behavioral observation (Fig. 4d) that lapses only on the manipulated side are affected. Like other models of inattention, it predicts that manipulating reward on one side should affect both lapses.

**Supplementary Figure 5:**
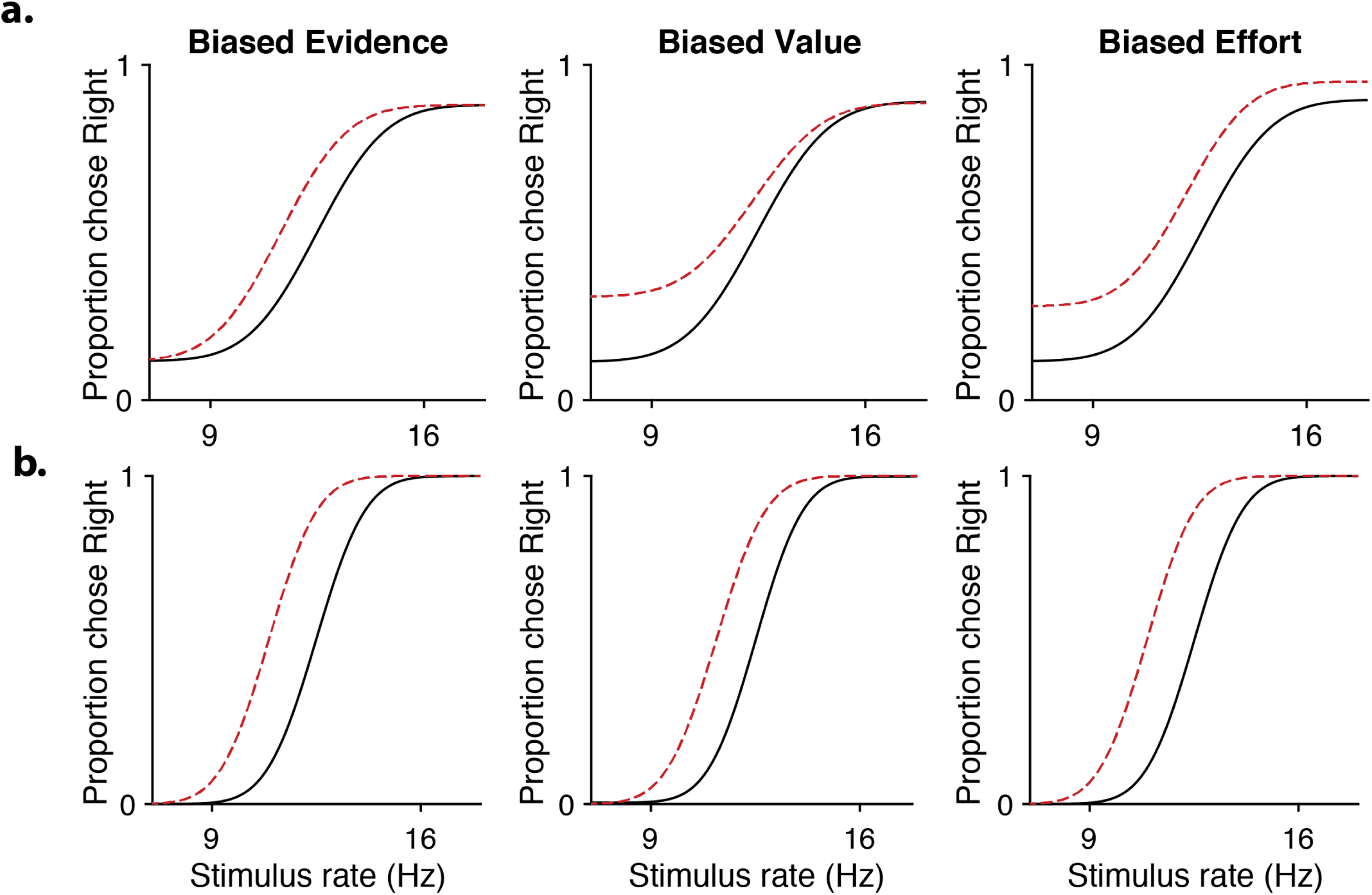
Lapses differentiate perturbations to different stages of the decision-making process. (a) Model predictions for biased sensory evidence (left), decreased contralateral action value (center) and increased effort in performing contralateral movements (right). The three kinds of perturbations affect decisions at the sensory, value or motor stages and predict different effects on lapses. (b) All three perturbations reduce to the same effect (horizontal shift) in the absence of lapses.

**Supplementary Figure 6:**
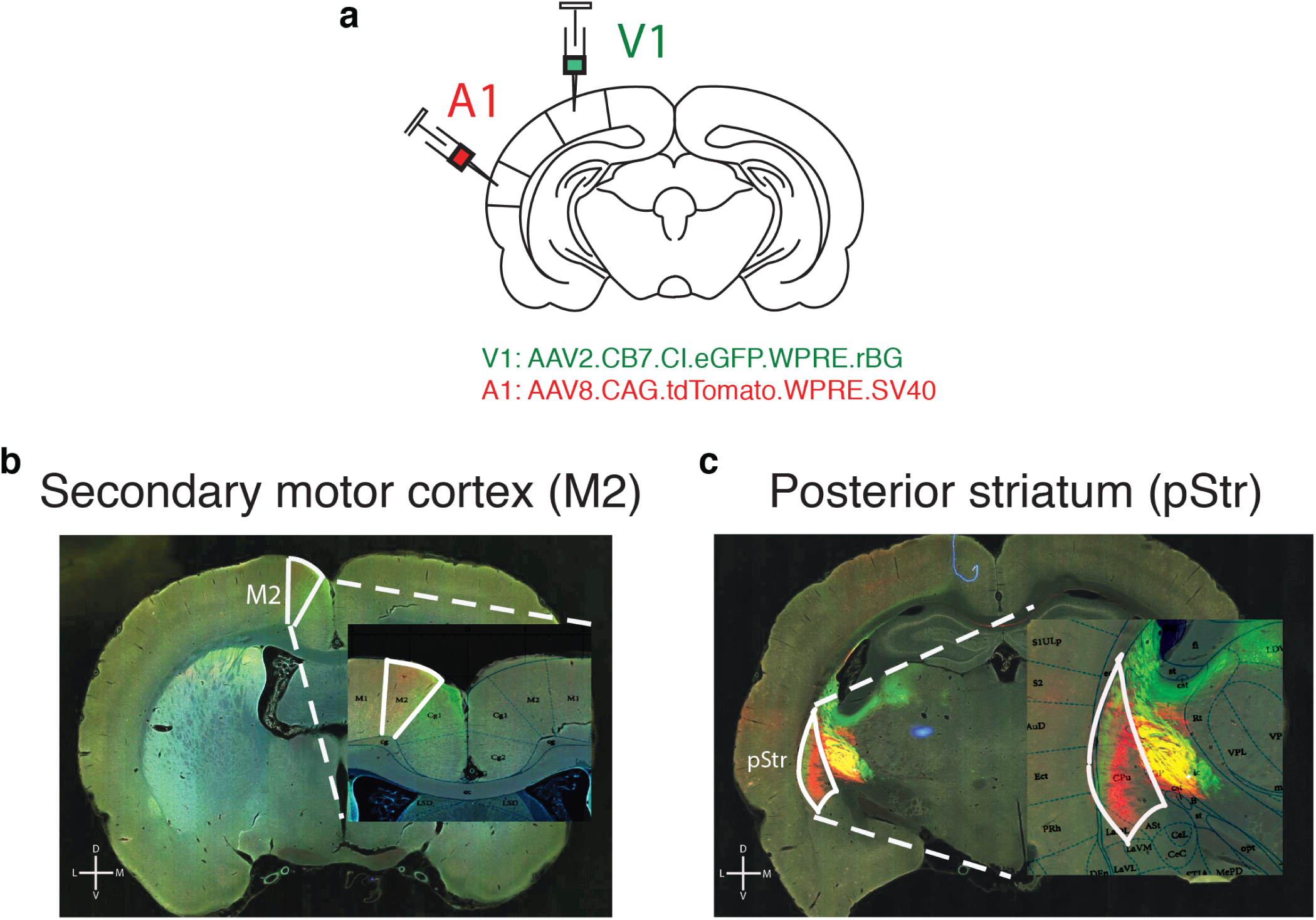
pStr and M2 receive direct projections from visual and auditory cortex. (a) Schematic of tracing experiments. AAV2.CB7.CI.EGFP.WPRE.RBG and AAV2.CAG.tdTomato.WPRE.SV40 constructs were injected unilaterally to primary visual (V1) and auditory (A1) cortices, respectively (V1 coordinates: 6.9 mm posterior to Bregma; 4.2 mm to the right of midline; A1 coordinates: 4.7 mm posterior to Bregma; 7 mm to the right of midline). (b) Secondary motor cortex (M2) receives inputs from V1 and A1 as shown by green and red fluorescence. (c) Posterior striatum (pStr) receives direct inputs from V1 and A1 as shown by green and red fluorescence. Yellow signal medial to pStr reflects overlapping passing fibers.

**Supplementary Figure 7:**
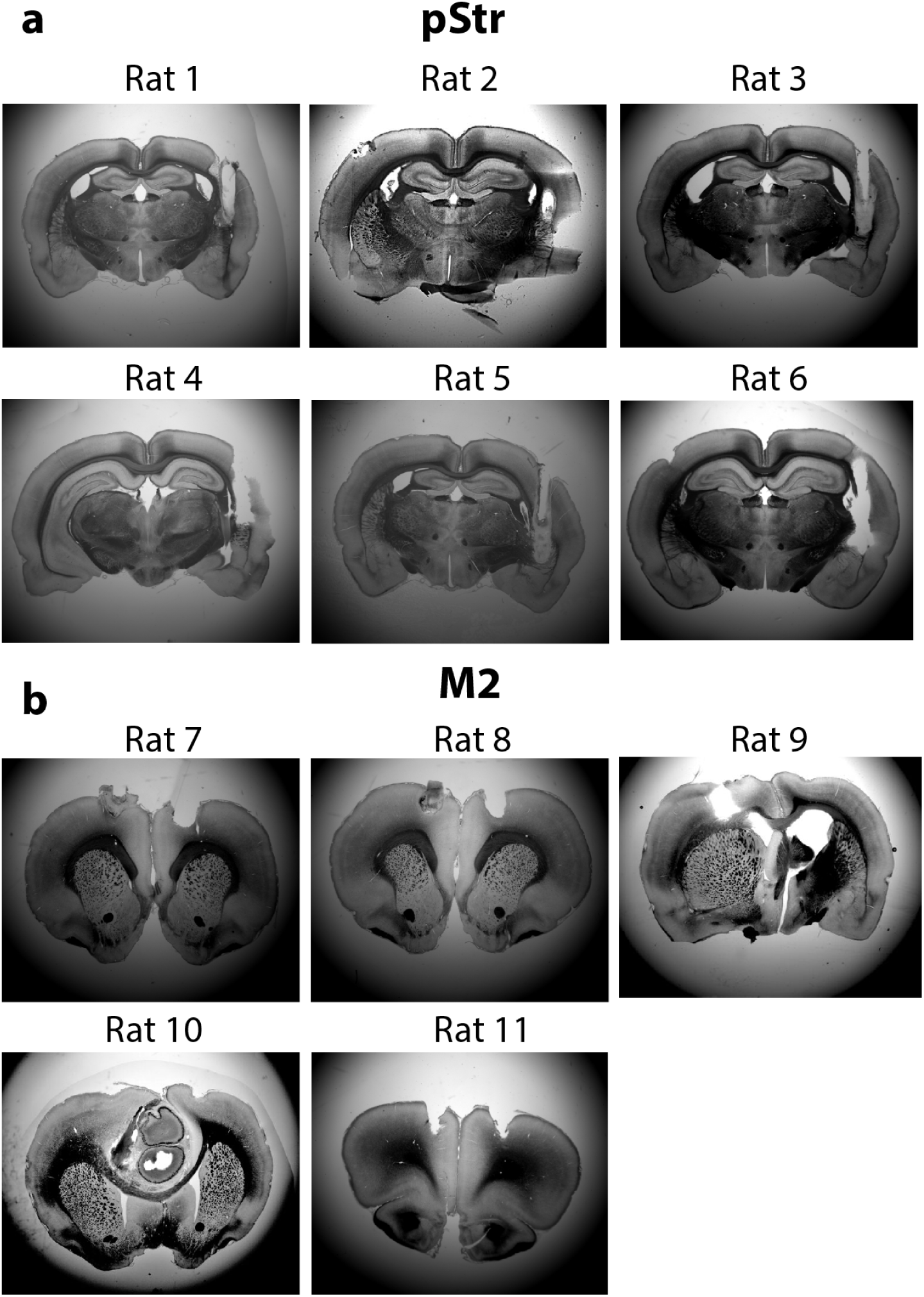
Histological slices of implanted rats. Representative coronal slices of all rats implanted with cannulae for muscimol inactivation experiments. (a) 6 rats were bilaterally implanted in posterior striatum (pStr). (b) 5 rats were implanted in secondary motor cortex (M2).

**Supplementary Figure 8:**
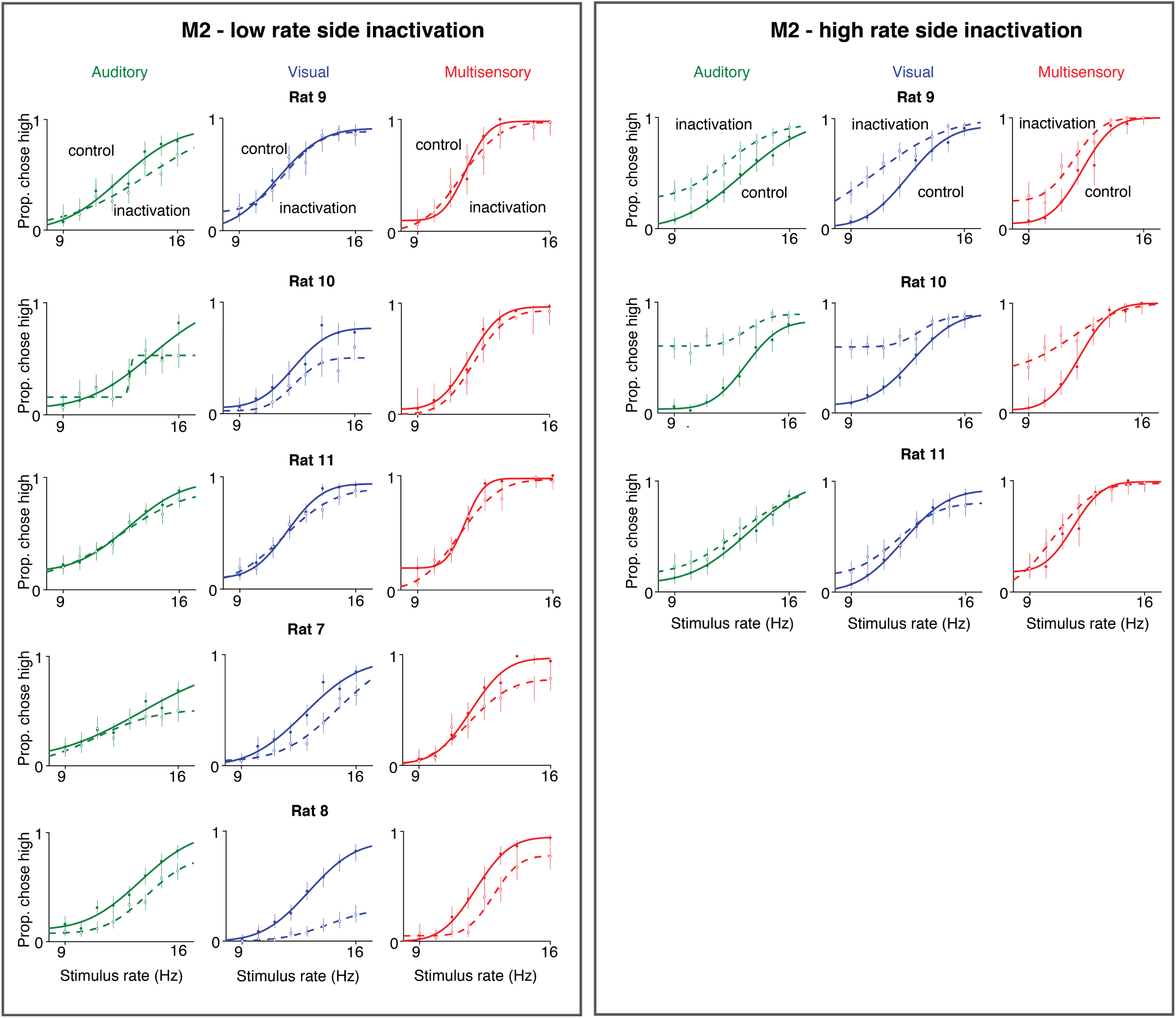
Single rat performance following M2 inactivation. Left: inactivation of the low-rate associated side. Rat shows increased lapses on high-rate trials on all sensory modalities. Right: inactivation of the high-rate associated side. Rat shows increased lapses on low-rate trials on all sensory modalities. Auditory (green), visual (blue) and multisensory (red).

**Supplementary Figure 9:**
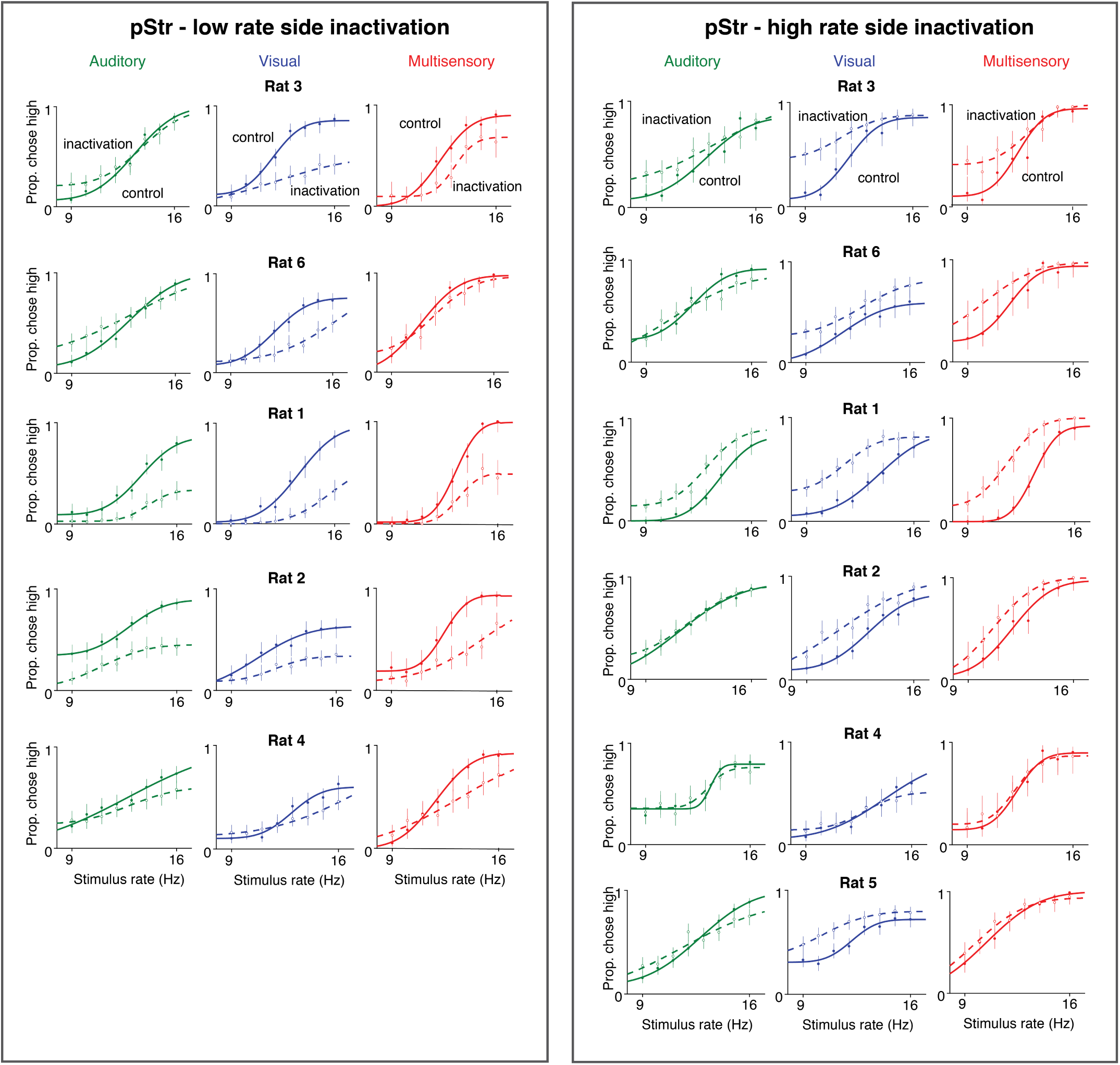
Single rat performance following pStr inactivation. Left: inactivation of the low-rate associated side. Rat shows increased lapses on high-rate trials on all sensory modalities. Right: inactivation of the high-rate associated side. Rat shows increased lapses on low-rate trials on all sensory modalities. Auditory (green), visual (blue) and multisensory (red).

**Supplementary Figure 10:**
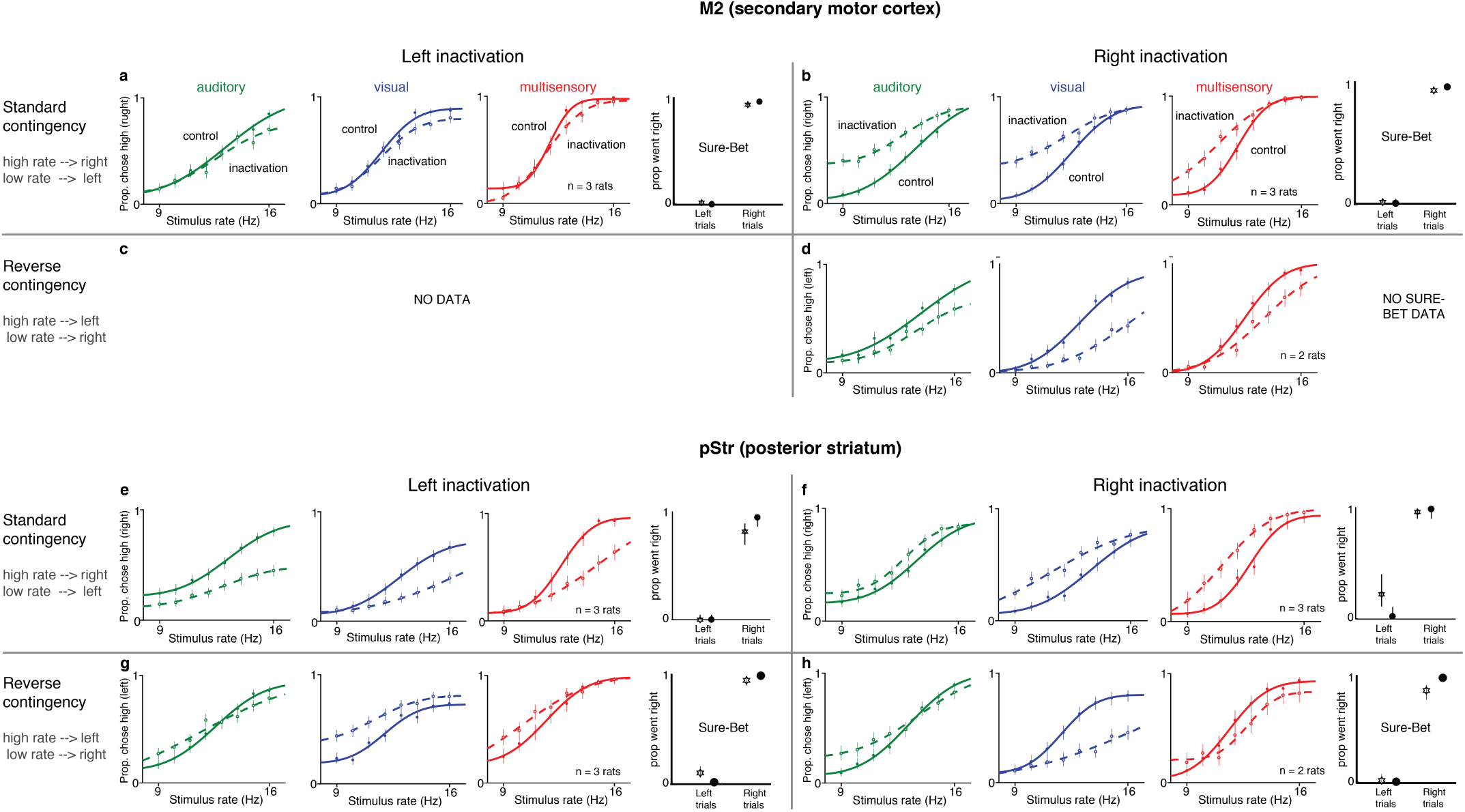
Unilateral inactivation of M2 or pStr biases performance ipsilaterally and increases contralateral lapses. Performance of the same rats shown in Figure 5b depicted as a function of the inactivated side (right or left) and the rate-contingency in which they were trained (standard or reverse). Standard contingency: high rate = go right, low rate = go left; reverse contingency: high rate = go left, low rate = go right. Each quadrant shows 4 plots: 3 psychometrics for rate discrimination trials and one for performance on sure-bet trials. auditory (green), visual (blue) and multisensory (red). (a)-(d) M2 inactivation. (e)-(h) pStr inactivation. (a), (d) Rats trained on the standard contingency and inactivated on the left hemisphere show increased lapses on the high rates (i.e., fewer rightward choices on high rates). No effect on sure-bet trials. (b), (f) Rats trained on the standard contingency and inactivated on the right hemisphere show increased lapses on the low rates (i.e., fewer leftward choices on low rates). No effect on sure-bet trials. (c), (g) Rats trained on the reverse contingency and inactivated on the left hemisphere show increased lapses on the low rates (i.e., fewer rightward choices on low rates). No effect on sure-bet trials. No data for this condition for M2 inactivation. (d), (h) Rats trained on the reverse contingency and inactivated on the right hemisphere show increased lapses on the high rates (i.e., fewer leftward choices on high rates). No effect on sure-bet trials for pStr inactivated animals; no data for M2 inactivated animals.

**Supplementary Figure 11:**
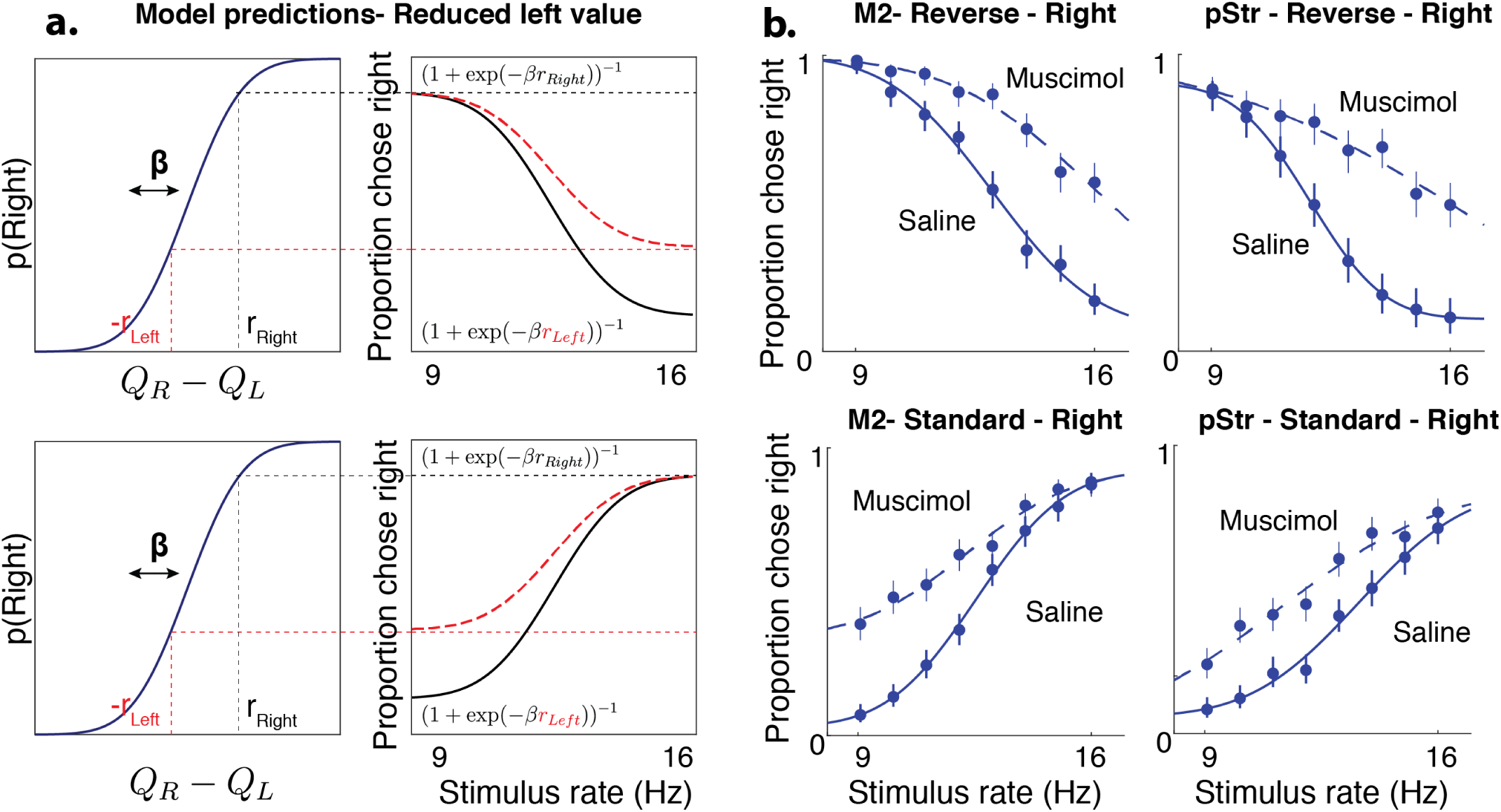
Inactivations devalue contralateral actions irrespective of associated stimulus. (a) Model predictions for rightward inactivations on standard (top) and reversed (bottom) stimulus-response contingencies - in both cases, the model predicts that reduced leftward action values should only affect lapses on the side associated with leftward movements. (b) Inactivation data on visual trials from M2 (left) or pStr (Right) shows a pattern of effects consistent with action value deficits, irrespective of the contingency.

**Supplementary Figure 12:**
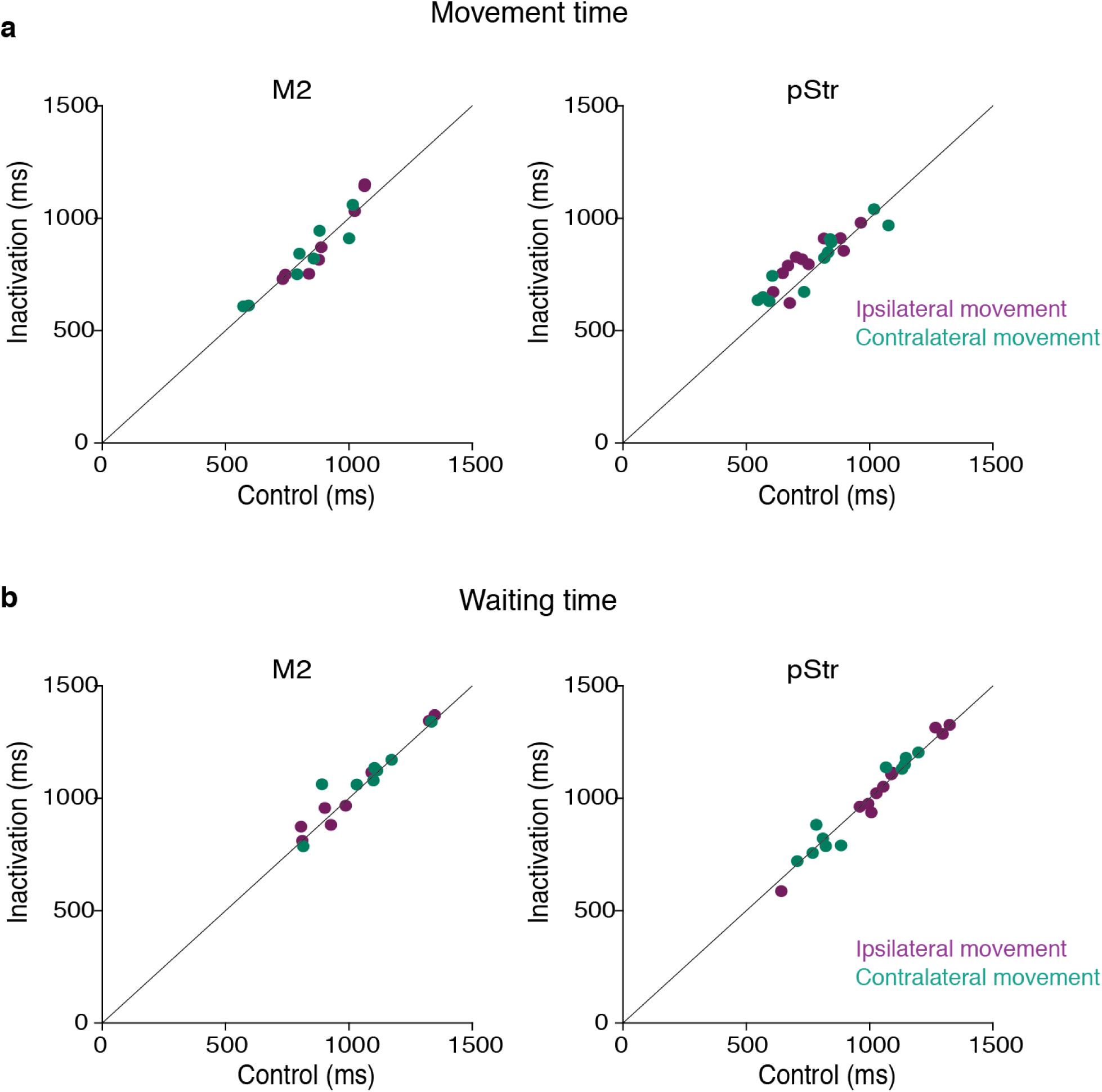
No significant effect on movement parameters following muscimol inactivation. (a) Mean movement times from the center port to the side ports were not significantly different following muscimol inactivation of M2 (left; p = 0.9554 for contralateral, 0.9852 for ipsilateral movements; n=5 rats) or pStr (right; p = 0.6629 for contra, p =0.2615 for ipsi, n=6 rats). Control data on the abscissa is plotted against inactivation data on the ordinate. Purple, movement toward the side ipsilateral to the inactivation site; blue, movement toward the side contralateral to the inactivation site; Error bars (s.e.m.) are not visible because they were obscured by the markers in all cases. (b) Mean wait times in the center port were not significantly different following muscimol inactivation of M2 (left; p = 0.7612 for contra, p =0.8896 for ipsi, n=5 rats) or pStr (right; p = 0.9128 for contra, p =0.9412 for ipsi, n=6 rats). All p-values were computed from paired t-tests. Error bars (s.e.m.) are not visible because they were obscured by the markers in all cases.

